# Transmissible SARS-CoV-2 variants with resistance to clinical protease inhibitors

**DOI:** 10.1101/2022.08.07.503099

**Authors:** Seyed Arad Moghadasi, Emmanuel Heilmann, Ahmed Magdy Khalil, Christina Nnabuife, Fiona L. Kearns, Chengjin Ye, Sofia N. Moraes, Francesco Costacurta, Morgan A. Esler, Hideki Aihara, Dorothee von Laer, Luis Martinez-Sobrido, Timothy Palzkill, Rommie E. Amaro, Reuben S. Harris

**Affiliations:** Department of Biochemistry, Molecular Biology, and Biophysics, University of Minnesota – Twin Cities, Minneapolis, Minnesota, USA, 55455; Institute of Virology, Medical University of Innsbruck, Innsbruck, Austria; Texas Biomedical Research Institute, San Antonio, Texas, USA, 78227; Department of Zoonotic Diseases, Faculty of Veterinary Medicine, Zagazig University, Zagazig, Egypt, 44511; Department of Pharmacology and Chemical Biology, Baylor College of Medicine, Houston, Texas, USA, 77030; Department of Chemistry and Biochemistry, University of California San Diego, La Jolla, California, USA, 92093; Department of Biochemistry and Structural Biology, University of Texas Health San Antonio; San Antonio, Texas, USA, 78229; Howard Hughes Medical Institute, University of Texas Health San Antonio; San Antonio, Texas, USA, 78229

**Keywords:** antiviral drugs, natural genetic variation, Paxlovid (nirmatrelvir), Xocova (ensitrelvir), protease inhibitor resistance, SARS-CoV-2 main protease (M^pro^/3CL^pro^)

## Abstract

Vaccines and drugs have helped reduce disease severity and blunt the spread of SARS-CoV-2. However, ongoing virus transmission, continuous evolution, and increasing selective pressures have the potential to yield viral variants capable of resisting these interventions. Here, we investigate the susceptibility of natural variants of the main protease (M^pro^/3CL^pro^) of SARS-CoV-2 to protease inhibitors. Multiple single amino acid changes in M^pro^ confer resistance to nirmatrelvir (the active component of Paxlovid). An additional clinical-stage inhibitor, ensitrelvir (Xocova), shows a different resistance mutation profile. Importantly, phylogenetic analyses indicate that several of these resistant variants have pre-existed the introduction of these drugs into the human population and are capable of spreading. These results encourage the monitoring of resistance variants and the development of additional protease inhibitors and other antiviral drugs with different mechanisms of action and resistance profiles for combinatorial therapy.

**One Sentence Summary:** Resistance to protease inhibitor drugs, nirmatrelvir (Paxlovid) and ensitrelvir (Xocova), exists in SARS-CoV-2 variants in the human population.

## INTRODUCTION

The main protease (M^pro^/3CL^pro^) of coronaviruses (CoV) is an attractive target for drug development, initially pursued in response to the first severe acute respiratory syndrome (SARS) pandemic in 2002 and swiftly revisited in response to the more recent SARS-CoV-2 (SARS2)/COVID-19 pandemic (*1-4*). M^pro^ activity is essential for virus replication and, combined with precedents set by the successful development of human immunodeficiency virus-type 1 (HIV-1) and hepatitis C virus (HCV) protease inhibitors, drugs targeting this enzyme are likely to help treat SARS2 infections (*5, 6*). Many groups have embarked on campaigns to target M^pro^ with multiple chemical series being advanced into potent inhibitors at unprecedented speeds, mainly owing to the wealth of biochemical and structural information that has accumulated on coronaviruses proteases over the past two decades (*7-11*). For instance, prior knowledge accelerated the development of PF-00835231 into PF-07321332 (nirmatrelvir), the active ingredient in Paxlovid and the first M^pro^ inhibitor to be used clinically (*12*). Another M^pro^ inhibitor that recently received emergency use authorization in Japan is S-217622 (ensitrelvir, Xocova), a non-covalent, non-peptidic inhibitor developed through computational and medicinal chemistry (*13, 14*). Ensitrelvir and other molecules in various stages of development may soon provide alternatives to Paxlovid and present opportunities for combinatorial therapy.

While Paxlovid is already proving useful in blunting SARS2 disease pathogenesis, the long-term consequences of wide-spread use are unknown. Resistance is a major concern given the relatively rapid rates at which SARS2 is changing (Alpha, Beta, Delta, Omicron, *etc*.) and the fact that the potency of nirmatrelvir and ensitrelvir varies widely against other coronavirus species (*15*). For instance, the main proteases of the human α-coronaviruses NL63 and 229E are less susceptible to these drugs suggesting that natural mechanisms of resistance may already exist in nature (*12, 16*). In addition, during the clinical development of Paxlovid, murine coronavirus MHV-A59 was used to study nirmatrelvir resistance (*17*). One of the selected amino acid changes, corresponding to S144A in SARS2, causes a >90-fold reduction in the binding efficacy (*K*_*i*_) of nirmatrelvir to recombinant M^pro^ *in vitro*.

We recently reported a cell-based gain-of-signal assay for SARS2 M^pro^ function in which wildtype (WT) protease activity suppresses luminescent signal by cleaving cellular substrates to prevent accumulation of reporter mRNA and, importantly, this suppressive effect can be overcome by genetic or chemical inhibition of M^pro^ to yield signal increases proportional to mutant severity or inhibitor efficacy, respectively (*16*). This system enabled us to show that single amino acid changes (P168S and P168G) in an active site-adjacent β-hairpin of M^pro^ improve susceptibility to the HCV protease inhibitor boceprevir, yet have no effect on the inhibitory capacity of the tool compound GC376 [(*16*); repeated below]. P168S is a naturally occurring variant that accounts for 76% of amino acid changes at this position in SARS2 clinical isolates based on sequences deposited in the GISAID database [1-July-2022; (*18*)]. This example of differential drug responsiveness inspired us to use evolution- and structure-guided approaches here to ask whether this natural variation at position 168, other natural changes at position 168, and other naturally occurring variants in the vicinity of the M^pro^ active site cavity may confer resistance to nirmatrelvir and/or ensitrelvir. Our results combine to demonstrate that multiple drug resistance mutations already exist in transmissible isolates of SARS2 in the global population. However, some M^pro^ variants that show resistance to nirmatrelvir still retain full susceptibility to ensitrelvir and *vice versa*, consistent with distinct mechanisms of action and the possibility that the latter drug can be used if the former fails and *vice versa*.

## RESULTS

### A natural M^pro^ variation ΔP168 confers resistance to nirmatrelvir and ensitrelvir

As introduced above, P168S is the most frequent amino acid change observed at this position in SARS2 M^pro^. The next most frequent change at this position in the GISAID database is a single residue deletion, ΔP168 (21% of changes at this position on 1-July-2022; **Fig. 1A**; **Table 1**). High resolution structures show that P168 is located close to the binding sites of boceprevir and nirmatrelvir (4.0□ and 3.3□, respectively), and it is approximately twice as far from that of ensitrelvir [8.8□; (*14, 19*)] (**Fig. 1B**). Based on our prior work with the P168S variant (*16*), we predicted that changes at this amino acid position in M^pro^ would more strongly affect the efficacy of boceprevir and nirmatrelvir and have little effect on ensitrelvir. Using our cell-based gain-of-signal assay (**Fig. S1A**), P168S causes a 5.5-fold increase in susceptibility to boceprevir in confirmation of our prior studies (**Fig. 1C**, left). We were therefore surprised to find that ΔP168 has no detectable effect with boceprevir, as its dose response curve is indistinguishable from that of WT M^pro^. In contrast, the ΔP168 variant shows 5.1- and 6.8-fold increased resistance to nirmatrelvir and ensitrelvir, respectively, whereas P168S variant maintains WT-like responsiveness to these two drugs (**Fig. 1C**, middle and right; **Table 1**). As a positive control, the S144A mutant described in the Introduction also shows a strong resistance phenotype for both nirmatrelvir and ensitrelvir with 12.2- and 16.9-fold increases in IC_50_ compared to WT M^pro^, respectively (**Fig. S1B**; **Table 1**).

**Table 1.**
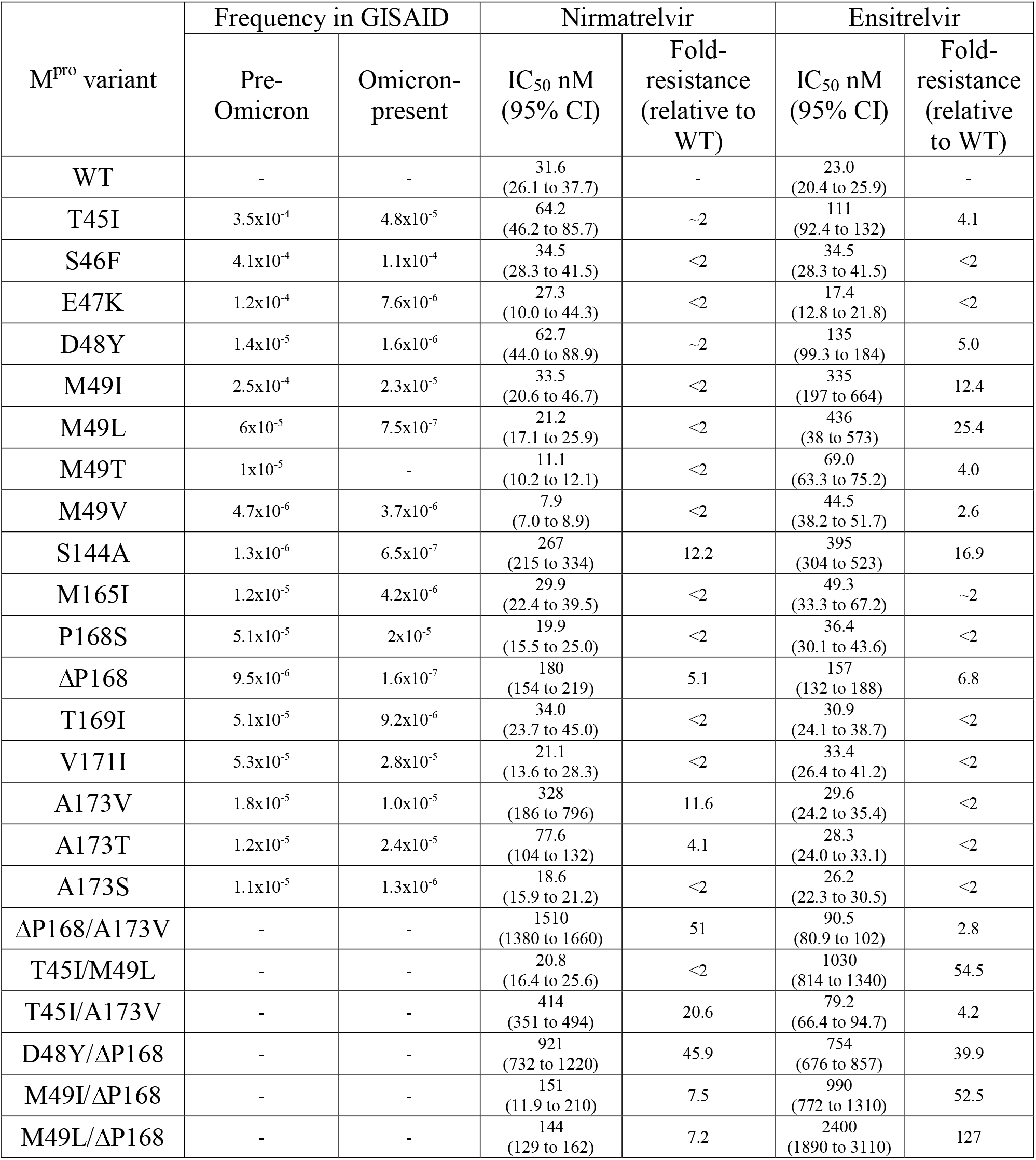
Summary of nirmatrelvir and ensitrelvir resistance phenotypes of SARS2 M^pro^ variants tested in the live cell gain-of-signal assay. Fold-resistance for each mutant tested was calculated based on relative IC_50_ in the live cell gain-of-signal assay versus WT in assays ran in parallel to limit potential variability due to transfection efficiency. Frequency of variants based on sequences deposited in the GISAID database as of 10-December-2022 (7,391,988 “Pre-Omicron” sequences and 6,830,473 “Omicron-present” sequences).

| M<sup>pro</sup> variant | Frequency in GISAID |  | Nirmatrelvir |  | Ensitrelvir |  |
| --- | --- | --- | --- | --- | --- | --- |
|  | Pre-Omicron | Omicron-present | IC<sub>50</sub> nM (95% CI) | Fold-resistance (relative to WT) | IC<sub>50</sub> nM (95% CI) | Fold-resistance (relative to WT) |
| WT | - | - | 31.6<br>(26.1 to 37.7) | - | 23.0<br>(20.4 to 25.9) | - |
| T45I | 3.5x10<sup>-4</sup> | 4.8x10<sup>-5</sup> | 64.2<br>(46.2 to 85.7) | ~2 | 111<br>(92.4 to 132) | 4.1 |
| S46F | 4.1x10<sup>-4</sup> | 1.1x10<sup>-4</sup> | 34.5<br>(28.3 to 41.5) | <2 | 34.5<br>(28.3 to 41.5) | <2 |
| E47K | 1.2x10<sup>-4</sup> | 7.6x10<sup>-6</sup> | 27.3<br>(10.0 to 44.3) | <2 | 17.4<br>(12.8 to 21.8) | <2 |
| D48Y | 1.4x10<sup>-5</sup> | 1.6x10<sup>-6</sup> | 62.7<br>(44.0 to 88.9) | ~2 | 135<br>(99.3 to 184) | 5.0 |
| M49I | 2.5x10<sup>-4</sup> | 2.3x10<sup>-5</sup> | 33.5<br>(20.6 to 46.7) | <2 | 335<br>(197 to 664) | 12.4 |
| M49L | 6x10<sup>-5</sup> | 7.5x10<sup>-7</sup> | 21.2<br>(17.1 to 25.9) | <2 | 436<br>(38 to 573) | 25.4 |
| M49T | 1x10<sup>-5</sup> | - | 11.1<br>(10.2 to 12.1) | <2 | 69.0<br>(63.3 to 75.2) | 4.0 |
| M49V | 4.7x10<sup>-6</sup> | 3.7x10<sup>-6</sup> | 7.9<br>(7.0 to 8.9) | <2 | 44.5<br>(38.2 to 51.7) | 2.6 |
| S144A | 1.3x10<sup>-6</sup> | 6.5x10<sup>-7</sup> | 267<br>(215 to 334) | 12.2 | 395<br>(304 to 523) | 16.9 |
| M165I | 1.2x10<sup>-5</sup> | 4.2x10<sup>-6</sup> | 29.9<br>(22.4 to 39.5) | <2 | 49.3<br>(33.3 to 67.2) | ~2 |
| P168S | 5.1x10<sup>-5</sup> | 2x10<sup>-5</sup> | 19.9<br>(15.5 to 25.0) | <2 | 36.4<br>(30.1 to 43.6) | <2 |
| ΔP168 | 9.5x10<sup>-6</sup> | 1.6x10<sup>-7</sup> | 180<br>(154 to 219) | 5.1 | 157<br>(132 to 188) | 6.8 |
| T169I | 5.1x10<sup>-5</sup> | 9.2x10<sup>-6</sup> | 34.0<br>(23.7 to 45.0) | <2 | 30.9<br>(24.1 to 38.7) | <2 |
| V171I | 5.3x10<sup>-5</sup> | 2.8x10<sup>-5</sup> | 21.1<br>(13.6 to 28.3) | <2 | 33.4<br>(26.4 to 41.2) | <2 |
| A173V | 1.8x10<sup>-5</sup> | 1.0x10<sup>-5</sup> | 328<br>(186 to 796) | 11.6 | 29.6<br>(24.2 to 35.4) | <2 |
| A173T | 1.2x10<sup>-5</sup> | 2.4x10<sup>-5</sup> | 77.6<br>(104 to 132) | 4.1 | 28.3<br>(24.0 to 33.1) | <2 |
| A173S | 1.1x10<sup>-5</sup> | 1.3x10<sup>-6</sup> | 18.6<br>(15.9 to 21.2) | <2 | 26.2<br>(22.3 to 30.5) | <2 |
| ΔP168/A173V | - | - | 1510<br>(1380 to 1660) | 51 | 90.5<br>(80.9 to 102) | 2.8 |
| T45I/M49L | - | - | 20.8<br>(16.4 to 25.6) | <2 | 1030<br>(814 to 1340) | 54.5 |
| T45I/A173V | - | - | 414<br>(351 to 494) | 20.6 | 79.2<br>(66.4 to 94.7) | 4.2 |
| D48Y/ΔP168 | - | - | 921<br>(732 to 1220) | 45.9 | 754<br>(676 to 857) | 39.9 |
| M49I/ΔP168 | - | - | 151<br>(11.9 to 210) | 7.5 | 990<br>(772 to 1310) | 52.5 |
| M49L/ΔP168 | - | - | 144<br>(129 to 162) | 7.2 | 2400<br>(1890 to 3110) | 127 |

**Fig. 1.**
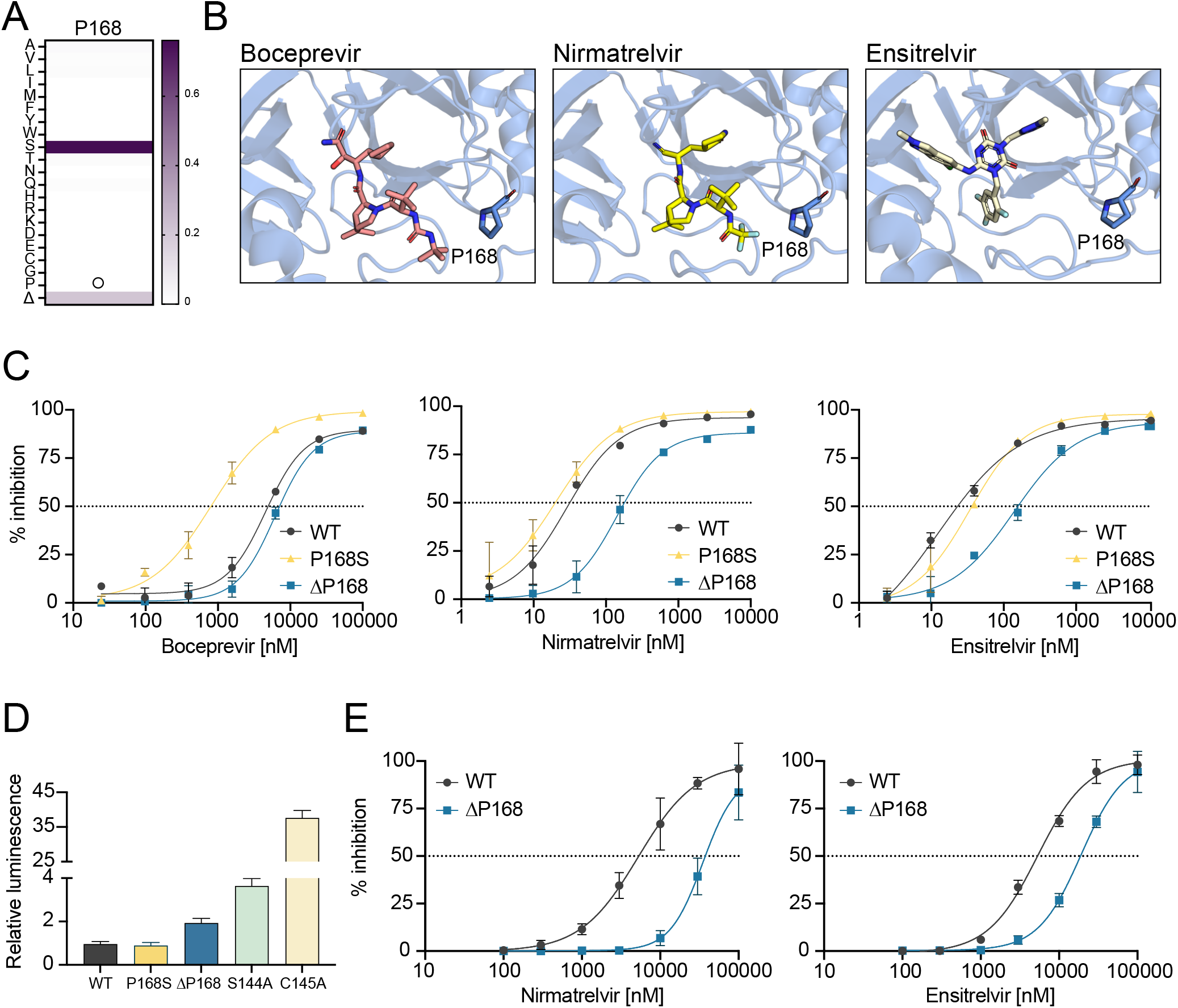
ΔP168 confers resistance to nirmatrelvir and ensitrelvir. **(A)** Relative frequency of amino acid changes at P168, excluding proline (open circle), in SARS2 genomes (1-July-2022, GISAID database). **(B)** Co-crystal structures of SARS2 M^pro^ in complex with boceprevir (PDB: 6WNP), nirmatrelvir (PDB: 7SI9), and ensitrelvir (PDB: 7VU6). **(C)** Dose-response curves of WT, P168S, and ΔP168 M^pro^ variants using the live cell Src-M^pro^-Tat-fLuc assay with 4-fold serial dilution of inhibitor beginning at 10 µM for nirmatrelvir and ensitrelvir or 100 µM for boceprevir (data are mean +/- SD of biologically independent triplicate experiments). **(D)** Relative luminescence of cells expressing Src-M^pro^-Tat-fLuc variants in the absence of inhibitor. **(E)** Dose-response curves of nirmatrelvir and ensitrelvir against WT and ΔP168 M^pro^ in an orthologous VSV-based M^pro^ *cis*-cleavage assay (data are mean +/- SD of biologically independent triplicate experiments).

An additional metric of our cell-based assay for M^pro^ function is background luminescent signal in the absence of a protease inhibitor (*16*). The WT construct yields very low luminescence and any diminution in M^pro^ catalytic activity results in increased signal with a maximum of approximately 40-fold as defined by a catalytic inactivating mutation (C145A, **Fig. 1D**). Consistent with our previous biochemical work (*16*), P168S shows no change in background luminescence relative to WT, whereas the genetically selected mutant S144A causes a 3.5-fold increase (**Fig. 1D**), in line with a reported decrease in biochemical activity (*20*). In comparison, ΔP168 elicits an intermediate, 2-fold increase in background luminescence (**Fig. 1D**). These results suggest that ΔP168 M^pro^ has near-WT catalytic activity and may be capable of supporting virus replication (addressed directly below). In support of both inhibitor resistance and protease activity results, the increased resistance of the M^pro^ ΔP168 variant to both nirmatrelvir and ensitrelvir is also apparent using an orthogonal VSV-based M^pro^ *cis-*cleavage assay in which inhibition of catalytic activity enables VSV replication (*21*) (**Fig. 1E**; **Table 2**).

**Table 2:** Resistance phenotypes of M^pro^ variants using the VSV-based *cis*-cleavage system. Fold-resistance is calculated by relative IC_50_ versus WT in assays ran in parallel (*, asterisk indicates an estimated value due to highest drug concentration failing to restore 50% activity).

| M<sup>pro</sup> variant | Nirmatrelvir |  | Ensitrelvir |  |
| --- | --- | --- | --- | --- |
|  | IC<sub>50</sub> μM<br>(95% CI) | Fold-resistance<br>(relative to<br>WT) | IC<sub>50</sub> μM<br>(95% CI) | Fold-resistance<br>(relative to<br>WT) |
| WT | 11.2<br>(10.8 to 11.7) | - | 12.9<br>(12.2 to 14.4) | - |
| T45I | 13.0<br>(11.2 to 15.4) | <2 | 48.1<br>(45.9 to 50.7) | 3.7 |
| D48Y | 15.7<br>(14.6 to 17.1) | <2 | 27.5<br>(26.0 to 28.7) | 2.1 |
| M49I | 4.1<br>(3.8 to 4.4) | <2 | >100* | >10* |
| M49L | 6.7<br>(6.5 to 6.9) | <2 | >100* | >10* |
| ΔP168 | 35.5<br>(32.2 to 47.2) | 7.1 | 19.1<br>(16.9 to 20.1) | 3.5 |
| A173T | 45.3<br>(40.0 to 55.2) | 4.0 | 18.8<br>(18.3 to 19.3) | <2 |
| A173V | 82.6<br>(59.5 to 93.5) | 7.4 | 10.3<br>(3.8 to 27.3) | <2 |
| ΔP168/A173V | >100* | >10* | 37.4<br>(33.2 to 43.0) | 3.1 |
| T45I/ΔP168 | >100* | >10* | >100* | >10* |
| T45I/A173V | >100* | >10* | 27.4<br>(24.2 to 29.6) | 2.2 |
| D48Y/ΔP168 | >100* | >10* | >100* | >10* |
| M49I/ΔP168 | 33.1<br>(32.0 to 34.2) | 3.0 | >100* | >10* |

### Drug resistance profiles of additional naturally occurring single amino acid M^pro^ variants

Encouraged by the resistance phenotypes caused by amino acid changes at a single position, we extended our analyses to include nine additional naturally variable residues that localize to two separate regions in proximity to the M^pro^ active site (**Fig. 2A-B**; **Table 1**). First, given the strong phenotypes observed with P168 variants, we hypothesized that mutations at additional residues within this β-hairpin might also confer drug resistance. Specifically, residues 165, 169, 171, and 173 show variability across coronavirus species (**Fig. 2A**) and, importantly, also within circulating SARS2 variants (**Fig. 2C**). M165, T169, and V171 were each substituted with isoleucine because this is a recurrent change at these positions, and A173 was substituted to valine as the most observed change at this position in SARS2 and also the residue found naturally in the human α-coronaviruses HCoV-229E and -NL63, both of which exhibit decreased susceptibility to nirmatrelvir (*12, 16*). The second region of interest encompasses M^pro^ residues 45-49, a small helix that forms the lid of the hydrophobic S2 subsite through the sidechain of M49. Notably, this is the most variable region across different coronaviruses species both in amino acid identity as well as overall length (**Fig. 2A**). For instance, M49I is a frequent change in circulating viruses (>1000 occurrences) and a substitution we have previously shown to have little effect in GC376 or boceprevir susceptibility (*16*), but its impact on nirmatrelvir and ensitrelvir efficacy has yet to be analyzed. Furthermore, we were intrigued by the hydrogen bonding pattern formed by T45 and D48 and curious whether naturally occurring changes that disrupt these interactions (T45I and D48Y) might destabilize the helical structure and impact inhibitor binding (**Fig. 2B**). Similarly, S46F and E47K were selected as non-conservative changes that may also disrupt the secondary structure of this region and impact inhibitor responsiveness.

**Fig. 2.**
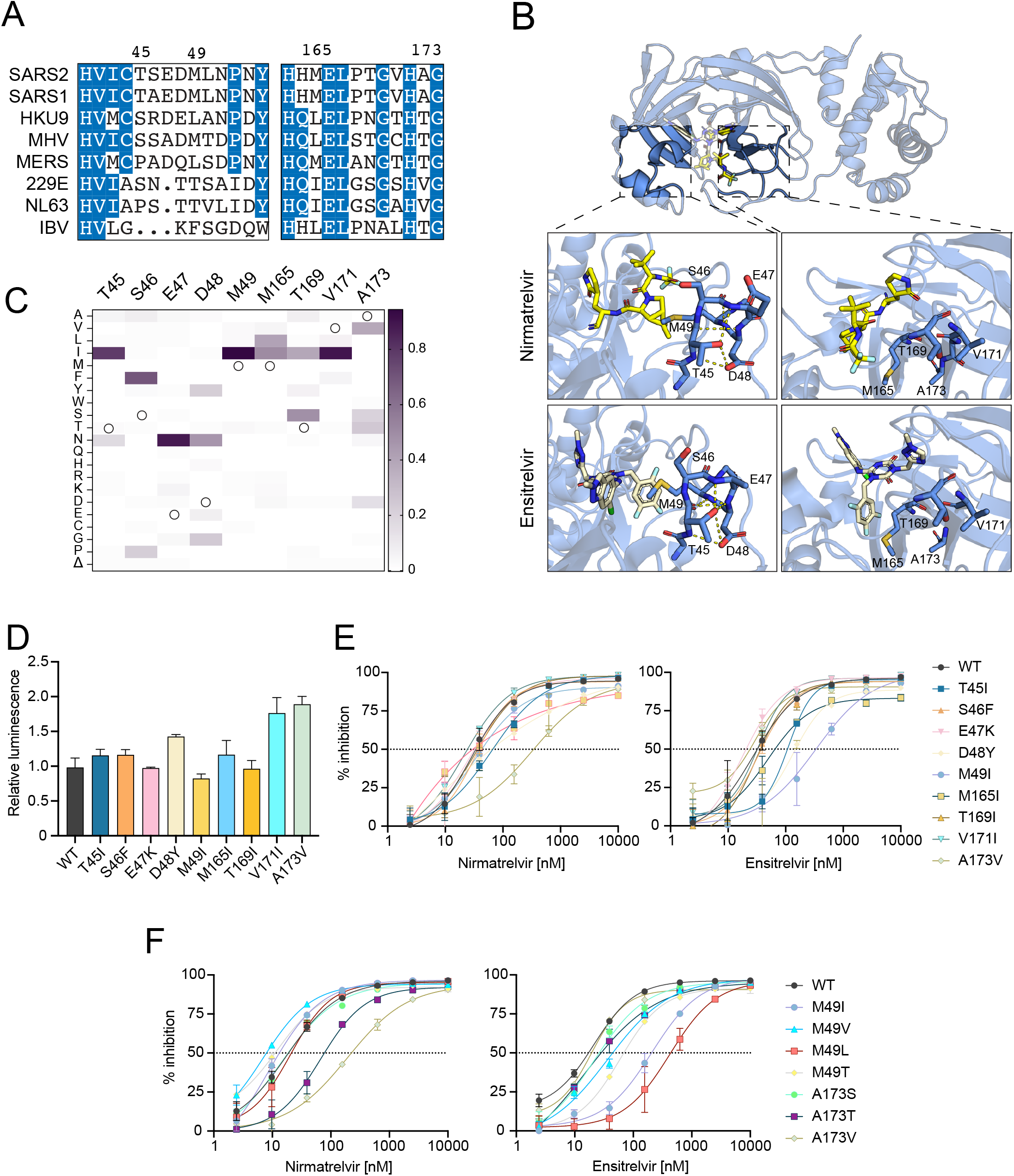
Variable active site residues elicit differential resistance to nirmatrelvir and ensitrelvir. **(A)** Alignment of coronavirus M^pro^ amino acid sequences spanning residues 41-54 and 163-174 (based on SARS2 M^pro^ residue position). **(B)** Structure of SARS2 M^pro^ and inhibitors highlighting variable residues that can be mutated (PDB: 7SI9 and 7VU6 for nirmatrelvir and ensitrelvir, respectively). **(C)** Relative frequency of amino acid changes at tested variable residues, excluding the respective WT residue (open circle), in SARS2 genomes (1-July-2022, GISAID database). **(D)** Relative luminescence of cells expressing Src-M^pro^-Tat-fLuc variants in the absence of inhibitor. (**E-F**) Dose-response curves of variants using the live cell Src-M^pro^-Tat-fLuc assay with 4-fold serial dilution of inhibitor beginning at 10 µM (data are mean +/- SD of biologically independent triplicate experiments).

Of the nine amino acid substitution mutants described above, five (S46F, E47K, M165I, T169I, and V171I) have little effect (<2-fold) on M^pro^ susceptibility to either nirmatrelvir or ensitrelvir and cause no substantial increases in background luminescence, consistent with near-WT catalytic activity (**Fig. 2D-E**; **Table 1**). In contrast, A173V immediately stands out as a separation-of-function variant by causing a 11.6-fold increase in resistance to nirmatrelvir and no change in susceptibility to ensitrelvir (**Fig. 2E**; **Table 1**). Importantly, A173V does not appear to drastically impact protease activity given the modest, 2-fold increase in background luminescence in the absence of inhibitor compared to WT M^pro^ (**Fig. 2D**). Additional contrast is seen with mutations surrounding the S2 subsite. T45I and D48Y cause 4.1- and 5-fold increases in resistance to ensitrelvir relative to the WT, and both have more modest effects on nirmatrelvir susceptibility (approximately 2-fold; **Fig. 2E**; **Table 1**). Strikingly, M49I causes no shift in the nirmatrelvir dose response in comparison to a 12.4-fold increase in resistance to ensitrelvir (**Fig. 2E**; **Table 1**).

Given the contrasting effects on inhibitor resistance caused by A173V and M49I, additional amino acid substitutions were generated for these two positions to gain additional insights into resistance mechanisms. A173T and A173S are variants observed in GISAID at frequencies similar to A173V; however, A173S causes no resistance to nirmatrelvir, whereas A173T causes an intermediate 4.1-fold resistance phenotype, suggesting that the bulkiness of the side chain of residue 173 correlates with the magnitude of nirmatrelvir resistance (**Fig. 2F**; **Table 1**). Similar to A173V, A173T and A173S do not affect susceptibility to ensitrelvir (**Fig. 2F**). M49V, M49T, and M49L are also found in circulating isolates, but at lower frequencies than M49I (**Table 1**). M49V and M49T cause milder phenotypes with 2.6- and 4-fold resistance to ensitrelvir, respectively (**Fig. 2F**; **Table 1**). In contrast, M49L causes 25.4-fold resistance to ensitrelvir and no change in nirmatrelvir susceptibility (**Fig. 2F**; **Table 1**). Notably, MERS has a leucine at the equivalent position in M^pro^ (**Fig. 2A**), and it exhibits higher ensitrelvir antiviral EC_50_ values compared to SARS and SARS2 (*14*). All these additional variants at M49 and A173 tested exhibit less than 2-fold changes in background luminescence compared to WT M^pro^ suggesting negligible changes in catalytic activity (**Fig. S2**). All single amino acid changes that exhibit resistance phenotypes using the gain-of-signal assay show similar results with nirmatrelvir and ensitrelvir in an orthogonal VSV-based M^pro^ assay (*21*) (**Fig. S3**; **Table 2**).

### A double mutant of M^pro^ with synergistic resistance to nirmatrelvir

Within our panel of naturally occurring single amino acid M^pro^ variants, two of the largest effects on nirmatrelvir resistance are ΔP168 and A173V (5.1- and 11.6-fold, respectively). This prompted us to test whether the combination might be additive or multiplicative in terms of drug resistance. Remarkably, the ΔP168/A173V double mutant shows a 51-fold increase in resistance to nirmatrelvir (**Fig. 3A**; **Table 1**). In contrast, this double mutant elicits only a 2.8-fold increase in resistance to ensitrelvir, which is less than that of the ΔP168 mutant alone (compare response curves in **Fig. 3A** and **Fig. 1C**, and numeric values in **Table 1**). As above, these single mutations have modest effects on background luminescence levels relative to WT, and the double mutant elicits a roughly additive effect with less than 3-fold increase in overall luminescence indicative of protease functionality (**Fig. 3B**). The VSV-based system yields similar resistance phenotypes for the single and double mutants (**Table 2**). These results combine to suggest that a strong resistance to nirmatrelvir can be achieved by combining two naturally occurring amino acid changes.

**Fig. 3.**
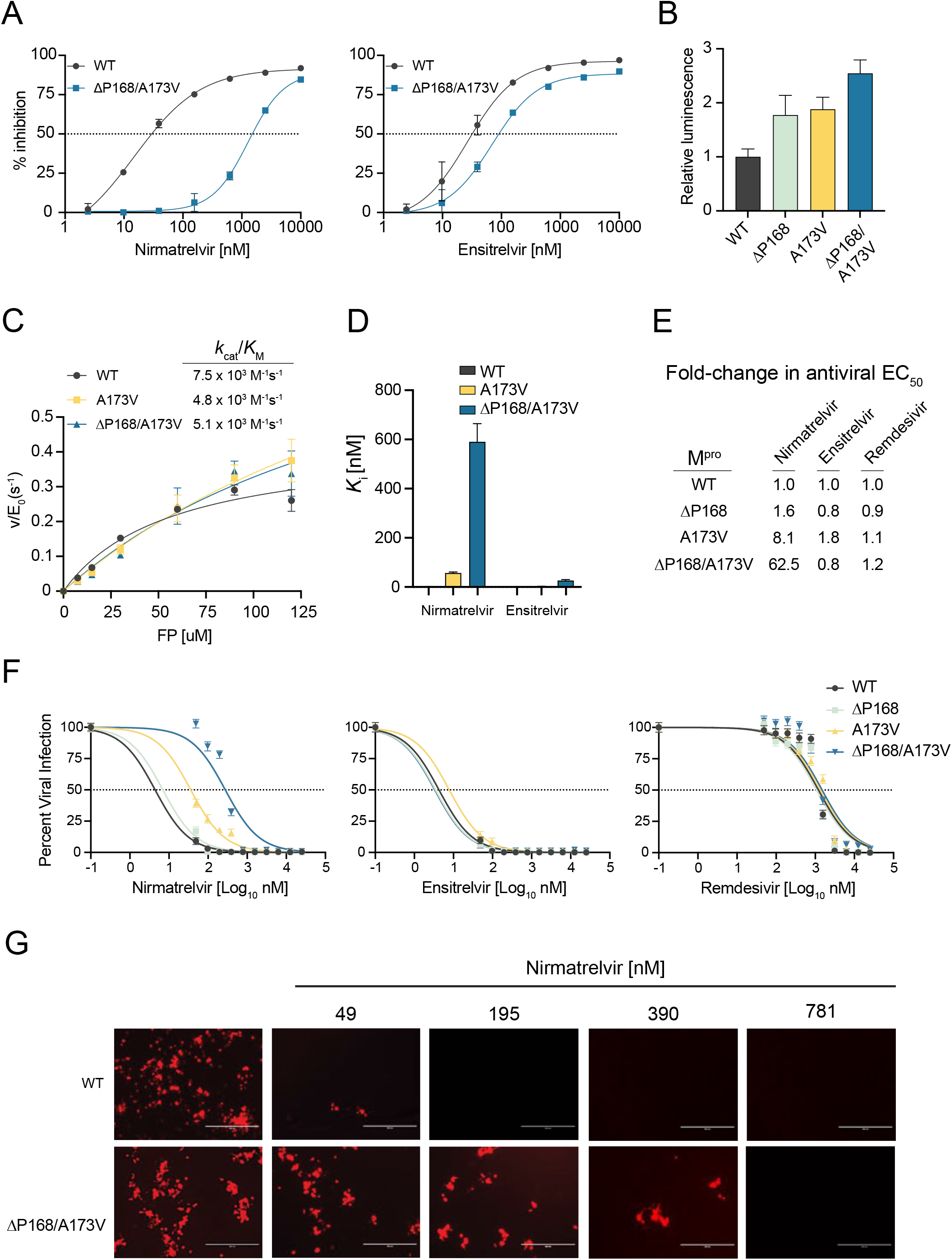
ΔP168/A173V double mutant elicits synergistic selective resistance to nirmatrelvir. **(A)** Dose-response of ΔP168/A173V mutant vs WT using the live cell Src-M^pro^-Tat-fLuc assay with 4-fold serial dilution of inhibitor beginning at 10 µM. **(B)** Relative luminescence of cells expressing respective Src-M^pro^-Tat-fLuc variants in the absence of inhibitor. **(C)** Kinetic parameters of purified M^pro^ variants *in vitro* using the Dabcyl-KTSAVLQSGFRKM-E(Edans)-NH_2_ FRET peptide (FP) as a substrate. **(D)** *K*_*i*_ of nirmatrelvir and ensitrelvir for purified M^pro^ variants derived using the Morrison equation with kinetic parameters calculated from data in panel C. (**E-F**) Antiviral activity of nirmatrelvir, ensitrelvir, and remdesivir with the indicated recombinant SARS2 viruses in A549-hACE2 cells (2-fold dilution series beginning at 25 µM, data are mean +/- SD of biologically independent quadruplicate experiments). (**G**) Representative fluorescence microscopy images of mCherry-expressing WT and ΔP168/A173V SARS2 infections following dosage with the indicated concentrations of nirmatrelvir.

To directly characterize the biochemical properties of these mutants, recombinant M^pro^ was purified from *E. coli* with an N-terminal SUMO-tag and a C-terminal His-tag, which are removed during purification to generate enzymes with authentic N- and C-termini (*22*). The WT, A173V, and ΔP168/A173V enzymes purify to near homogeneity but, despite multiple attempts, the ΔP168 single mutant was not amenable to purification [precise reason(s) unclear but potentially due to poor solubility and/or aggregation propensity in bacteria]. Michaelis-Menten kinetics for WT M^pro^ hydrolysis of a FRET peptide Dabcyl-KTSAVLQSGFRKM-E(Edans)-NH_2_ yield a *k*_cat_ of 0.43 s^-1^ and *k*_cat_/*K*_M_ of 7.5 × 10^3^ M^-1^s^-1^, consistent with prior values (*23, 24*) (**Fig. 3C**; **Fig. S4A**). The A173V substitution causes a 3-fold increase in *k*_cat_ and a less than 2-fold decrease in *k*_cat_/*K*_M_ (**Fig 3C**; **Fig. S4A**). The ΔP168/A173V enzyme also displays near-WT kinetic parameters with a 2-fold increased *k*_cat_ and a less than 2-fold decrease in *k*_cat_/*K*_M_ (**Fig. 3C**; **Fig. S4A**). These results indicate that neither the A173V nor the ΔP168/A173V enzyme exhibits a major decline in M^pro^ catalytic activity.

The same FRET-based system was then used to quantify inhibition by nirmatrelvir and ensitrelvir. WT M^pro^ is inhibited potently by nirmatrelvir with a *K*_i_ of 1.1 ± 0.95 nM, again consistent with prior values (*12*) (**Fig. 3D**; **Fig. S4B**). In contrast, nirmatrelvir is 50-fold less potent against A173V M^pro^ with a *K*_i_ of 57 ± 4.2 nM (**Fig. 3D**; **Fig. S4B**). Strikingly, nirmatrelvir is nearly 600-fold less potent against the ΔP168/A173V enzyme with a *K*_i_ value of 590 ± 74 nM (**Fig. 3D**; **Fig. S4B**). In comparison, ensitrelvir inhibits WT and A173V enzymes similarly with *K*_i_ values of 0.2 ± 0.56 nM and 2.3 ± 0.94 nM (*p* value = 0.19 by unpaired *t*-test). The *K*_i_ of ensitrelvir for the double mutant is higher (23 ± 4.1 nM) and, by deduction, this is likely due to ΔP168. These biochemical findings agree with results from the two live cell assays described above and collectively indicate that the A173V and ΔP168/A173V enzymes are active and that these naturally occurring amino acid changes confer a strong and preferential resistance to nirmatrelvir.

As an additional functional test, we built these resistance mutants of M^pro^ into a bacterial artificial chromosome (BAC)-based reverse genetics system (WA-1 strain), produced viral stocks using Vero-E6 cells, and performed a series of drug titration experiments as described (*25*). This system has an mCherry reporter linked to the viral N-gene and separated at the translation level by a 2A self-cleavage site, enabling quantification of virus infectivity and replication by fluorescence microscopy (*25*) (**Fig. S5A**). To determine drug susceptibility, A549-hACE2 cells were infected with virus encoding WT, ΔP168, A173V, or ΔP168/A173V M^pro^ at equal multiplicities of infection, then treated with varying concentrations of antiviral inhibitors, and virus replication was quantified 72 hours post-infection (hpi) by analyzing mCherry fluorescence. Whereas the virus encoding ΔP168 M^pro^ does not exhibit resistance to nirmatrelvir, A173V M^pro^ causes an 8.1-fold increase in EC_50_ and, strikingly, ΔP168 synergizes with A173V to cause a 62.5-fold increase in EC_50_ (**Fig. 3E-G**; **Fig. S5B**; **Table 3**). In contrast, none of these mutants alter virus susceptibility to ensitrelvir, which suggests that the level of resistance caused by ΔP168 alone in cell-based assays M^pro^ inhibition assays is insufficient to alter susceptibility in a viral context. Importantly, these results indicate that the ΔP168/A173V double mutant virus is still inhibited effectively by ensitrelvir, despite a strong resistance to nirmatrelvir (**Fig. 3E-F**). As expected, none of the mutants alter susceptibility to remdesivir, which acts by inhibiting the viral RNA dependent RNA polymerase (**Fig. 3E-F**). Surprisingly, whereas the ΔP168 virus shows WT-like replication kinetics, the A173V virus exhibits a spreading replication defect which remained unaltered when combined with ΔP168 (**Fig. S5C**). Although the catalytic efficiency of the A173V mutant is similar to the WT enzyme using a model substrate *in vitro* that mimics the Nsp4-5 cleavage site (above), it is possible that one or more natural cleavage sites in the viral polyprotein may be disproportionately affected by this change during authentic virus replication.

**Table 3:** Resistance phenotypes of M^pro^ variants in replication-competent SARS-CoV-2.

| M<sup>pro</sup> variant | Nirmatrelvir | Ensitrelvir | Remdesivir |
| --- | --- | --- | --- |
|  | IC<sub>50</sub> nM<br>(95% CI) | IC<sub>50</sub> nM<br>(95% CI) | IC<sub>50</sub> μM<br>(95% CI) |
| WT | 4.4<br>(3.4 to 5.5) | 4.3<br>(2.9 to 5.8) | 1.2<br>(0.59 to 2.5) |
| ΔP168 | 7.0<br>(4.1 to 10.2) | 3.6<br>(2.7 to 4.4) | 1.1<br>(0.59 to 2.0) |
| A173V | 36.0<br>(29.8 to 43.0) | 7.8<br>(6.2 to 9.5) | 1.3<br>(0.85 to 2.1) |
| ΔP168/A173V | 275<br>(161 to 465) | 3.3<br>(2.1 to 4.8) | 1.5<br>(0.78 to 2.9) |

### Possible structural explanations for key resistance phenotypes

To determine a structural basis for the differential resistance to nirmatrelvir and ensitrelvir, we performed molecular dynamics (MD) simulations and calculated root-mean square fluctuations (RMSF) to estimate per-residue perturbations related to ΔP168 and A173V. First, RMSF analyses of WT and variant M^pro^ enzymes trajectories indicate two regions of high flexibility – residues 40-65 and 185-195. Interestingly, the former region shows increased flexibility attributable to ΔP168 and A173V, whereas the latter region – like the rest of the protein – shows similar mobility regardless of variation (**Fig. 4A**; **Fig. S6**). Residues 40-65 comprise two α-helices that form a lid-like motif immediately above the S2 subsite of M^pro^ (top left in **Fig. 4B**). A partially unfolded state in the ΔP168 variant is observed for the α-helix directly adjacent to catalytic histidine (residues 43-53; top right in **Fig. 4B**), and this is exacerbated for the A173V and ΔP168/A173V variants where this region is frequently seen fully unfolded (bottom left and right, respectively, in **Fig. 4B**). Such open conformations may decrease capacity of M^pro^ for native substrate recognition and account for the higher observed *K*_*M*_ values (**Fig. S4**).

**Fig. 4.**
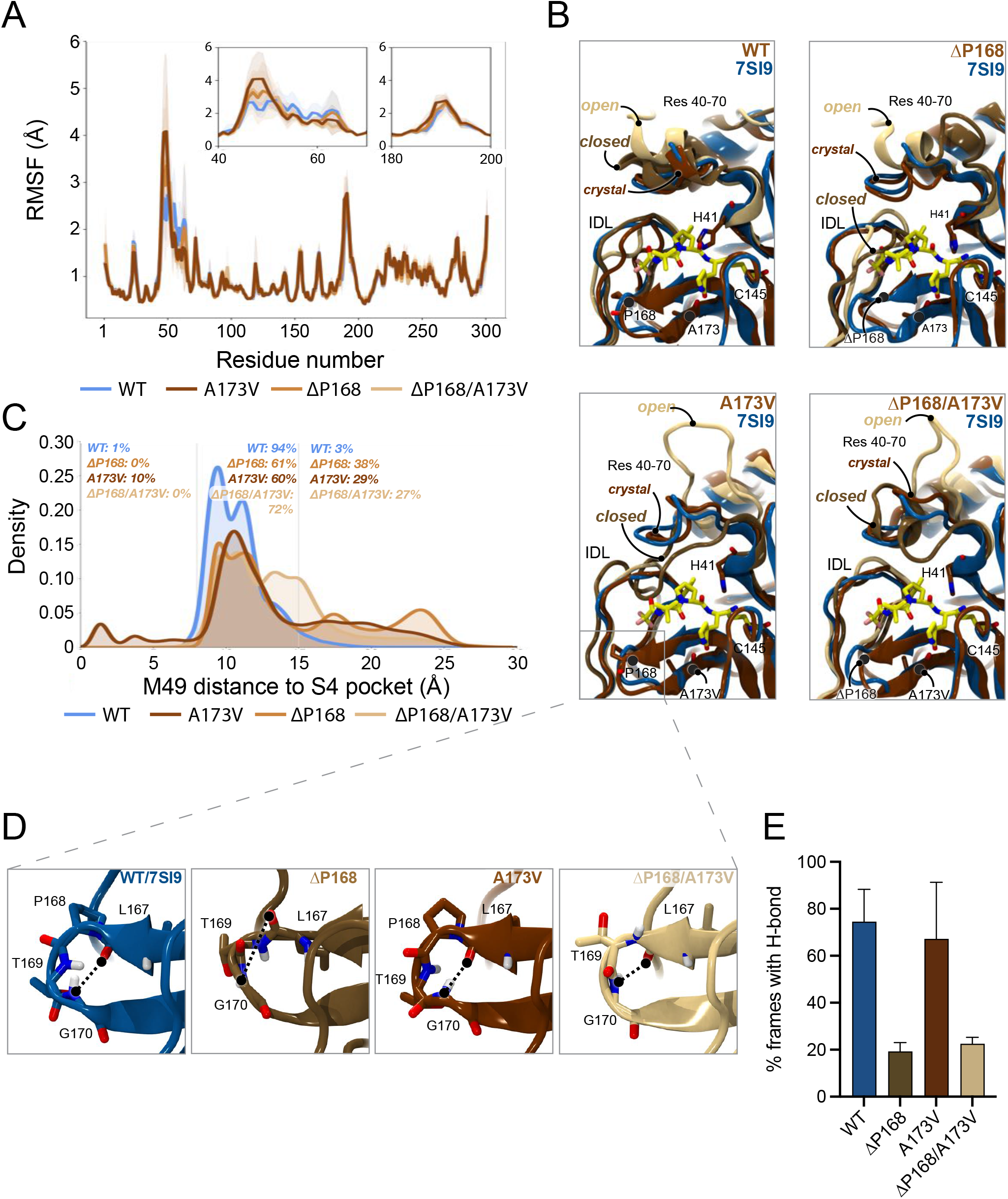
Structural interpretation of M^pro^ ΔP168/A173V synergistic resistance to nirmatrelvir. **(A)** Root-mean square fluctuations (RMSFs) calculated per residue for each simulated M^pro^ variant. To maintain numbering consistency, RMSFs for deleted residues are plotted as the interpolation of values from the prior residue (167) and the following residue (169). Calculated RMSFs per residue were averaged over all replicas and across chains A and Inset plots highlight increased flexibility of regions 40-70 and 180-200. **(B)** Molecular model images demonstrating the flexibility of residues 40-70 and 180-200 captured in simulations of WT, ΔP168, A173V, and ΔP168/A173V. All MD frames were aligned to the 7SI9 crystal structure (blue ribbons) to highlight the nirmatrelvir (carbon atoms represented in yellow licorice) binding site. MD simulation frames, in crystal, closed, and open conformations, are represented in chocolate brown, golden brown, and latte brown ribbons, respectively. **(C)** Histograms demonstrating the distribution of frames for which M49 penetrates the S4 subpocket and the distribution of frames for which M49 (and thus residues 43-53) moves far above M^pro^’s native binding grove. **(D)** Molecular model images demonstrating the binding site β-hairpin structure (residues 165-175) as seen in WT (blue ribbons), ΔP168 (golden brown), A173V (chocolate brown), and ΔP168/A173V (latte brown) MD frames. Hydrogen bonds, or lack thereof, between L167 backbone carbonyl and G170 backbone nitrogen are highlighted. **(E)** Percentage of frames calculated in which the L167-G170 backbone hydrogen bond is observed. Hydrogen bond requirements were established as: <4Å between L167 backbone carbonyl oxygen and G170 backbone nitrogen, and an angle of >120 degrees between L167 backbone carbonyl oxygen, G170 backbone hydrogen, and G170 backbone nitrogen (see results for complete characterization). Percentages were calculated and averaged per replica and per chain, and standard deviations used as error estimates.

In contrast, “closed” conformations of the same loop region may impart nirmatrelvir resistance. With A173V and ΔP168/A173V, the 43-53 loop encroaches upon the S4 subsite, a hydrophobic sub-pocket required for binding native substrates and the trifluoroacetamide moiety of nirmatrelvir (**Fig. 4B-C**; **Fig. S7A-C**). This is evidenced by increased variability in the distance between M49 and the S4 subsite (**Fig. 4C**). The closing of this loop region may also tighten the S2 subsite and clash with the fused cyclopropyl ring at the P2 position of nirmatrelvir (**Fig. 4B**). In comparison, ensitrelvir does not occupy either the S4 or S2 subsite and instead projects outwards into the S1’ subsite, which is consistent with our finding that A173V has little effect on the efficacy of ensitrelvir (**Fig. S7D**).

MD simulations also indicate that ΔP168 may negatively impact L167-G170 backbone H-bonding regardless of the residue (A or V) at position 173 (**Fig. 4D-E, Fig. S8**). The 165-175 β-hairpin sits above the interdomain loop (IDL, residues 180-200) to form the S3/4 subsite, and loss of this hydrogen bond has the potential to destabilize the hairpin structure. Notably, inward motion of the IDL is restricted by positioning of the 165-175 β-hairpin which is consistent with increased flexibility in the ΔP168 mutant (**Fig. 4A**). Whereas IDL structure modulates inhibitor binding, destabilization of the hairpin allows its encroachment into the S4 sub-pocket thereby decreasing inhibitor potency.

### Insights from additional double mutant combinations

Given the synergistic effect of ΔP168 and A173V on nirmatrelvir resistance, we next used our gain-of-signal cell-based assay to investigate double mutant combinations involving residues in the lid region described above (*i*.*e*., residues 40-70; **Fig. S9A-B**; **Table 1**). First, we combined the ensitrelvir resistant mutant T45I with single amino acid changes that confer resistance to ensitrelvir (M49L), nirmatrelvir (A173V), or both drugs (ΔP168). The T45I/M49L combination shows a synergistic 54.5-fold increase in ensitrelvir resistance, little change in susceptibility to nirmatrelvir, and no increase in background luminescence. The T45I/A173V combination exhibits a 20.6-fold increase in resistance to nirmatrelvir (more than additive), a 4.2-fold increase in resistance to ensitrelvir (near identical to T45I alone), and a modest 2-fold increase in background luminescence (similar to A173V alone). The effect of the T45I/ΔP168 combination could not be assessed accurately for drug resistance because it likely attenuates inferred M^pro^ catalytic activity as indicated by a >10-fold increase in background luminescence. This result is unexpected given that T45I and ΔP168 alone only modestly elevate background luminescence and suggests that this double mutant will not be infectious. Key double mutant results are recapitulated in the VSV-based M^pro^ cleavage assay (**Fig. S9C**; **Table 2**).

We also examined two double mutant combinations involving M49I, a mutation that alone confers 12-fold resistance to ensitrelvir and no change to nirmatrelvir (**Fig. S9A-B**; **Table 1**). M49I/ΔP168 and M49L/ΔP168 combinations confer a synergistic resistance to ensitrelvir of 52- and 127-fold, respectively, whereas resistance to nirmatrelvir is only approximately 7-fold for both double mutants (similar to ΔP168 alone). With regards to inferred catalytic activity, M49I/ΔP168 shows background luminescence indistinguishable from WT and M49L/ΔP168 has a 3.7-fold increase. These results highlight how even subtle changes in amino acid side chains can have significant effects on inhibitor resistance and/or enzyme catalysis. M49L double mutants were not tested in the VSV-based M^pro^ cleavage assay because this substitution alone already confers near-complete resistance to ensitrelvir (**Fig. S3**; **Table 1**).

### Global SARS2 variant distributions and evidence for transmission

The global frequency and distribution of an individual mutant indicates whether a particular amino acid change might be tolerated in nature. However, the current large sequence volume of SARS2 genomes in the GISAID database coupled with low/no stringency filtering results in the identification of mutations at every position of M^pro^, even at codons encoding conserved catalytic residues (**Fig. S10A**). This strongly suggests that the database contains a certain level of sequences that are not viable and therefore not transmitting through the human population. To address this issue, we used Ultrafast Sample placement on Existing tRee (UShER) to determine phylogenetic relationships between genomes harboring drug resistance mutations in M^pro^ (*26*). Importantly, analyzing mutational distances to common ancestors allows us to identify viral genomes that are hypermutated, such as a genome deposited recently in GISAID with >100 mutations relative to its closest ancestor, including M^pro^ C145A, which is likely a consequence of poor sequence coverage and annotation across the M^pro^ encoding region (**Fig. S10B-C**).

In comparison, viral genomes containing ΔP168 have arisen multiple times independently, with the majority occurring in the Delta lineage (**Fig. 5A**). Moreover, a distinct cluster of 49 genomes deposited between September and December of 2021 is derived from a single founder event followed by multiple regional transmissions in Germany, in addition to evidence consistent with spread to England, USA, Austria, and Romania (**Fig. 5B**; **Fig. S11**). Despite many independent occurrences of ΔP168 in prior lineages, only a single case has been documented in Omicron, identified in the Netherlands in November 2022 (**Fig. 5A**). This bias may result from the fact that all descendants of Omicron carry a characteristic P132H mutation in M^pro^ which might be functionally incompatible with ΔP168. In contrast, M49I and A173V have both occurred multiple times independently since the emergence of Omicron, with phylogenetic clusters indicative of transmission (**Fig. S11**). Overall, there is a higher frequency of M49I than M49L, which may reflect better relative fitness or simply be a consequence of the fact that three different single base substitutions can lead to an isoleucine codon and only two to a leucine (**Fig. 5C**). Moreover, similar to ΔP168, there is only a single introduction of M49L since the emergence of Omicron, which led to 4-related cases in Japan in early 2022 (**Fig. S11**). It is worth noting that these cases of M49L are of the BA 1.1.2 lineage which originated in Japan and contain an additional T169S change in M^pro^, which may provide a genetic background more tolerant to M49L or may constitute a compensatory mutation (**Fig. S11**).

**Fig. 5.**
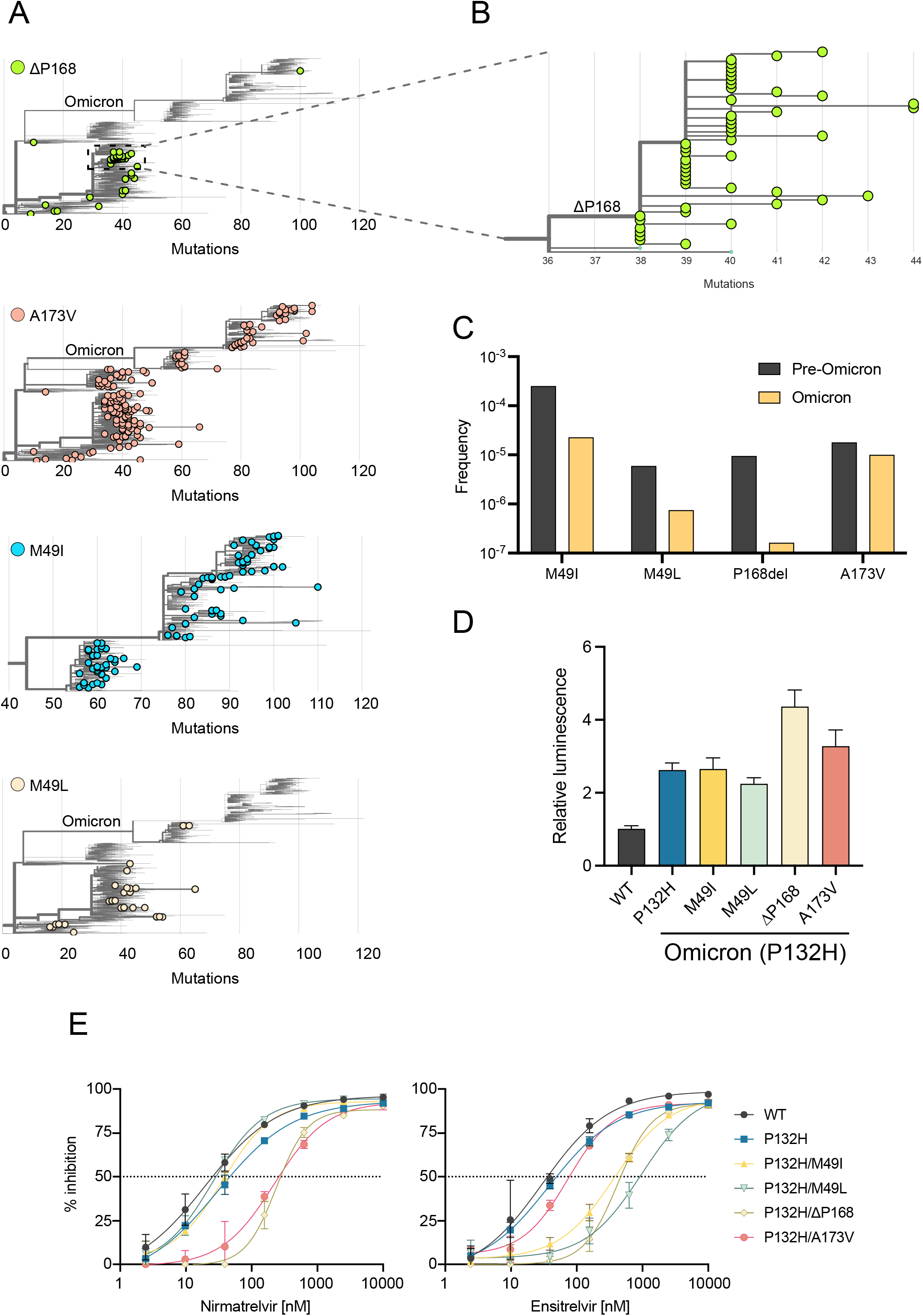
Phylogenetic analyses suggest mutual exclusivity for P132H and ΔP168. **(A)** Phylogenetic trees for viral genomes with the indicated resistance mutations (GISAID, 6-December-2022). Only the Omicron lineage is displayed for M49I due to the high frequency of this mutation. **(B)** Branches of the phylogenetic tree from panel A showing transmission of a Delta lineage isolate with ΔP168 likely from a single founder event. **(C)** Frequency of M^pro^ variants in Omicron lineage compared to all previous viral lineages. **(D)** Background luminescence as a proxy for protease activity of different M^pro^ variants on the P132H (Omicron) background using the live cell Src-M^pro^-Tat-fLuc assay. **(E)** Dose-response curves for nirmatrelvir and ensitrelvir inhibition of M^pro^ variants on the P132H (Omicron) background using the live cell Src-M^pro^-Tat-fLuc assay.

Given these apparent differences in the frequency of certain M^pro^ mutants since the emergence of Omicron, we asked whether the resistant phenotypes observed for M49I, M49L, ΔP168, and A173V could be recapitulated when combined with the P132H change in Omicron M^pro^. Using the live cell gain-of-signal assay, we observed that the P132H mutation alone causes a 2-fold increase in background luminescence relative to the ancestral WT sequence (**Fig. 5D**), which may be related to a decreased stability reported for this mutant enzyme *in vitro* (*25*). In combination with P132H, M49I and M49L do not cause additional increases in background luminescence, whereas ΔP168 and A173V both have an additional effect with the former being the most compromised (**Fig. 5D**). These results are consistent with phylogenetic observations above that P132H and ΔP168 may be functionally incompatible. However, most importantly, all mutants show similar drug resistance phenotypes when combined with P132H in our gain-of-signal assay, suggesting that these mutations are capable of conferring resistance to protease inhibitors in the context of currently circulating Omicron variants (**Fig. 5E**).

## DISCUSSION

Major efforts continue for developing antiviral drugs to complement vaccination-based strategies for treating patients infected by SARS2 with the ultimate hopes of ending the COVID-19 pandemic and fortifying against future coronavirus outbreaks. M^pro^ inhibitors are at the forefront of coronavirus antiviral drug development with Paxlovid (nirmatrelvir) already authorized for emergency clinical use in over 65 countries and several other compounds including ensitrelvir in various stages of development (*27*). However, drug resistance mutations have the potential to rapidly undermine these therapies. Here, we show that several naturally occurring M^pro^ variants already exhibit resistance to nirmatrelvir and ensitrelvir (results summarized in **Table 1** and **Table 2**). The highest levels of resistance resulting from a single amino acid substitution identified here is A173V for nirmatrelvir (11.6-fold) and M49L for ensitrelvir (25.4-fold). Phylogenetic analyses show that these (and other) variants have arisen multiple times independently in different parts of the globe with regional clusters and genetic linkage providing compelling evidence for transmission.

Nirmatrelvir is a substrate-mimicking covalent drug and ensitrelvir is a non-peptide/non-covalent inhibitor (*12, 14*). Consistent with distinct mechanisms of action, our studies indicate that these inhibitors are subject to at least partly non-overlapping resistance profiles. For instance, A173V confers selective resistance to nirmatrelvir, whereas M49I and M49L confer increased resistance to ensitrelvir. In comparison, ΔP168 appears to have a more broad-spectrum resistance phenotype. Encouragingly, antiviral assays with recombinant SARS2 corroborate our findings with two different cell-based assays and indicate that the ΔP168/A173V virus causes strong resistance to nirmatrelvir with little change in ensitrelvir susceptibility. Variation at additional residues may also produce distinct resistance patterns when present in isolation versus in combination. For instance, T45I and D48Y exhibit a mild preferential resistance to ensitrelvir as single changes; however, when combined with ΔP168, D48Y shows strong resistance to both inhibitors and T45I cripples enzyme activity. Although our structural modeling and MD simulations provide plausible explanations for the A173V (+/- ΔP168) resistance phenotype, additional dedicated studies will be needed to establish other mechanisms of action.

During revision of this manuscript, multiple preprints reported M^pro^ mutants with resistance to nirmatrelvir (*28-32*). A173V was selected during serial passage experiments in the presence of boceprevir and independently in the presence of nirmatrelvir, which coupled with our results suggest that A173 may be a resistance hot spot (*29, 30*). Our biochemical data using the purified A173V mutant demonstrate a lower affinity for nirmatrelvir as evidenced by a 50-fold increase in *K*_i_ with little change in catalytic efficiency against the canonical Nsp4-5 cleavage site. While these results support the selection of these mutants in serial passage, there appears to be some discrepancy in the magnitude of resistance between changes in *in vitro Ki* compared to changes in antiviral EC_50_. This is observed for other M^pro^ variants such as S144A, which is selected in serial passage and has been determined by Pfizer as causing a 90-fold increase in niramtrelvir *K*_*i*_ while antiviral studies show a more modest 2-fold increase in antiviral EC_50_ (*30*). The smaller changes in antiviral EC_50_ may result from the fact that M^pro^ has a wide range of affinities (*K*_*d*_ measurements ranging from 28 µM to 2.7 mM) for its different polyprotein cleavage sites (*33*). Therefore, while inhibitor affinity may be reduced by these mutants it could still be sufficient to bind the enzyme before cleavage of the lower affinity viral substrates. Therefore, resistance mutations may need to confer *K*_*i*_ increases of multiple orders of magnitude to cause large shifts in antiviral EC_50_. This interpretation is supported by our results with the ΔP168 mutant, which alone does not confer a shift in antiviral EC_50_, however ΔP168/A173V has 7.6-fold increase in EC_50_ compared to A173V alone (62.5-fold compared to WT). Our data with these select mutants are concordant across four orthologous assays (two live cell assays, *in vitro* biochemistry, and *in cellulo* with replication-competent virus) suggesting that multiple mutations may be necessary to decrease drug binding affinity and cause resistance.

Consistent with our gain-of-signal assay showing only a 2-fold increase in background luminescence, our kinetic analyses of the purified A173V mutant indicate similar catalytic efficiency to WT (<2-fold change in *k*_cat_/*K*_M_). However, the multicycle growth kinetic assays with recombinant SARS2 show decreased replication. Independent studies have reported no change in replication kinetics for A173V, whereas another saw a decrease similar to ours that could be rescued by an additional L50F mutation (*29, 30*). The reason for this discrepancy between replicative fitness of A173V from different labs is currently unclear, but different measurements of quantifying viral replication could be a contributing factor. Although activity is retained on the canonical Nsp4-5 substrate that is a standard for *in vitro* experiments, other cleavage sites along the polyprotein may be disproportionately impacted by this change. For instance, our MD simulations show an increase in the dynamics of amino acids 40-65, which form the top of the S2 subsite that accommodates the hydrophobic P2 position of the peptide substrate. As most cleavage sites along the viral polyprotein have a leucine at P2, phenylalanine and valine are also found at Nsp5-6 and Nsp6-7 junctions, respectively, which may be less efficiently processed by the A173V mutant. Importantly, however, this variant has sufficient activity for virus replication and is observed recurrently in patient sequences and, therefore, it has the potential to contribute to clinical resistance phenotypes.

Although many groups have focused appropriately on resistance to nirmatrelvir given its early emergency use authorization by the FDA, ensitrelvir resistance is now equally important to understand given that emergency use authorization was granted in Japan on 22-November-2022. Along with this approval, documentation was released on serial passage experiments selecting for D48G, M49L, P52S, and S144A as resistance mutations (*34*). These results support our finding of M49L showing the largest resistance phenotype using the gain-of-signal assay and the VSV-based assay. Furthermore, the selection of D48G also substantiates our hypothesis that disrupting hydrogen bonds between T45 and D48 to destabilize the structure of the helix above the S2 subsite can contribute to ensitrelvir resistance (indicated by our data for T45I and D48Y). Another recent report has also identified M49I as conferring selective resistance to ensitrelvir and elegantly demonstrates the structural basis of this phenotype being due to the bulky isoleucine reorienting H41 and disrupting a base stacking interaction with the inhibitor (*35*). This is consistent with our finding that M49L causes greater resistance than M49I due to branching of the leucine sidechain at the gamma carbon which is closer to H41. Together, these findings indicate that the 45-49 region of M^pro^ has the potential to become a hotspot for the development of ensitrelvir resistance mutations.

By using our facile live cell gain-of-signal assay coupled with sequence- and structure-informed mutation identification, we have been able to identify multiple changes in M^pro^ that confer varying degrees of resistance to nirmatrelvir and/or ensitrelvir. The resistance phenotypes described here are consistent between the four different assays we have implemented, and they are also consistent with reports by other groups through serial passage of virus or *in vitro* biochemical assays. Our cell-based gain-of-signal assay has the advantage of only requiring the transfection of a single plasmid, which increases the throughput of variant testing compared to generation of recombinant virus or purification of mutant enzymes (especially those that are difficult to purify such as the ΔP168 mutant). Using variants found within patient sequences at residues that are not strictly conserved across coronavirus species has helped identify changes more likely to be compatible with productive viral infection. However, it is important to take great care when classifying variants within the GISAID database as many annotated variants are likely to be sequence artefacts. For example, we found >4000 sequences with a M165Y change, and manual inspection revealed that this is due to a single guanosine deletion in a poly-U stretch, which causes a frameshift after F160 and leads simultaneously to “detection” of H163W, E166Q, and a downstream stop codon (**Fig. S12A-B**). Most of these sequences were produced using long-read nanopore technology which has a 76-fold higher rate of indel errors compared to short-read technology (*36*). These sequencing and mis-annotation mistakes can lead to incorrect conclusions regarding the presence of variants in the population and therefore manual inspection of viral genomes is encouraged for putative changes at strictly conserved residues or those requiring multiple base changes (*28*). Determining the phylogenetic relationships of different variants using publicly available tools such as UShER also provides additional validation of the emergence of these variants by identifying common ancestors and dynastic relationships within distinct geographic regions (*26*).

Although our gain-of-signal system provides robust metrics for evaluating mutants and M^pro^ inhibitors in live cells, there are also a few limitations. For instance, although the relative background luminescence can serve as a proxy for catalytic activity, it is currently unclear which cellular substrates are being cleaved to cause low reporter expression, thereby limiting the correlations that may be made to activity of M^pro^ against individual viral polyprotein cleavage sites. Moreover, our current approach focuses on amino acid residues that are variable (and not completely conserved) to avoid changes that would be severely deleterious which limits the potential number of resistance pathways tested. For instance, it is difficult to predict secondary suppressor mutations that could restore fitness of a resistant but deleterious mutation. An example of this is the identification of E166V and E166A selected during serial passage to confer a high level of resistance against nirmatrelvir and other peptidomimetic inhibitors (*29-31*). As E166 is a highly conserved residue, it was not tested here. Indeed, severe replications defects are evident with single substitutions at E166 but secondary mutations such as L50F and T21I appear to restore fitness (*29-31*). Thus, the gain-of-signal assay should be considered a valuable tool to study resistance mutations and complement traditional approaches such as serial passaging and competition studies to triage variants of interest that may be selected in cell culture or emerge *in vivo* during patient treatment before proceeding with more experimentally demanding approaches such as protein biochemistry and/or BSL3 testing with infectious viruses. Furthermore, we anticipate that the panel of mutants described here will be able to serve as an asset during the development of future generation M^pro^ inhibitors for rapid resistance profiling in parallel with structure activity relationship studies.

It is presently unclear what magnitude of resistance will be necessary for treatment failure in a clinical setting. Precedents with HCV NS3/4A show that single amino acid changes can elicit selective resistance of multiple orders of magnitude towards different inhibitors with minimal impact on viral fitness (*37*). However, resistance to HIV protease inhibitors often requires two or more mutations, with single amino acid changes typically showing modest changes in inhibitor susceptibility (*38, 39*). The naturally occurring SARS2 M^pro^ variants described here may serve as evolutionary stepping-stones for intermediate levels of resistance and provide a permissive environment enabling selection of secondary mutations that confer full drug resistance. Genetic surveillance of several of the variants identified here may be advantageous and strategies should be taken to minimize the widespread development of resistance including the careful design of M^pro^ inhibitor drugs with different resistance profiles, which encouragingly is likely to be the case for nirmatrelvir and ensitrelvir.

## MATERIALS AND METHODS

### Cell culture and M^pro^ reporter assays

The pcDNA5/TO-Src-M^pro^-Tat-fLuc reporter construct has been described (*16*). M^pro^ variants were generated by site-directed mutagenesis (primers available upon request), and all mutations were confirmed by Sanger sequencing. 293T cells were maintained at 37°C and 5% CO_2_ in DMEM (Gibco catalog number 11875093) supplemented with 10% fetal bovine serum (ThermoFisher catalog number 11965084) and penicillin-streptomycin (Gibco catalog number 15140122). For each M^pro^ variant, 3×10^6^ 293T cells were plated in a 10cm dish and transfected 24h later with 2µg of the corresponding Src-M^pro^-Tat-fLuc plasmid using TransIT-LT1(Mirus catalog number MIR 2304) transfection reagent. 4h post transfection, cells were washed once with phosphate buffered saline (PBS), trypsinized, resuspended in fresh media, counted, and subsequently diluted to a final concentration of 4×10^5^ cells/ml. Dilution series of inhibitors were prepared in fresh media at twice the final desired concentration of the reaction and 50µL was pipetted into a 96 well cell culture plate. 50µL of the cell suspension was added directly to the 96 well plate with inhibitor containing media to yield a final cell concentration of 20,000 cells per well. 44h after plating into 96 well plates, media was removed and 50µL of Bright-Glo reagent was added to each well and incubated at room temperature in the dark for 5m before transferring to white flat 96-well plate for measuring luminescence on a Biotek Synergy H1 plate reader. Percent inhibition at each concentration of inhibitor was derived with the formula below using the relative luminescence (RL) of an inhibitor treated sample to the untreated control.

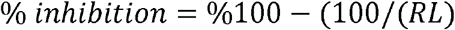

Results were plotted using GraphPad Prism 9 and fit using a four-parameter non-linear regression to calculate IC_50_. Resistance of mutants was calculated by the fold change in IC_50_ of the mutant relative to WT M^pro^.

The *cis*-cleaving VSV based M^pro^ assay was performed as described (*21*). Briefly, 293T cells were transfected with the phosphoprotein (P)-M^pro^ fusion construct for each variant of interest and after an overnight incubation, resuspended and plated into 96 well plates. Transfected cells were then treated with the intended inhibitor and infected with VSV-ΔP-RFP at an MOI of 0.1. 48h post-infection, fluorescence was measured using a Fluoro/ImmunoSpot counter (CTL Europe GmbH, Bonn, Germany). Data were plotted as relative fluorescence using GraphPad Prism 9 and fit using a four-parameter non-linear regression to calculate IC_50_.

### Protein expression and purification

SARS2 M^pro^ and mutants were expressed and purified from *E. coli* BL21(DE3) using a pSUMO-SARS2-M^pro^ protein expression plasmid and purification protocol described previously with minor variations (*22*). Briefly, M^pro^, A173V, and ΔP168/A173V mutants were expressed with an N-terminal SUMO-tag and a C-terminal His-tag. The M^pro^ open reading frame sequence was flanked on the N terminus by its endogenous cleavage site (SAVLQ↓SGFRK) and on its C terminus by a PreScission protease cleavage site (SGVTFQ↓GP). The N-terminal SUMO tag was cleaved by M^pro^ during protein expression in *E. coli*. Cells containing the pSUMO-SARS2-M^pro^ plasmid were grown in Luria broth (LB) at 37°C to an optical density 600 nm of 0.8 to 1 and M^pro^ expression was induced with 0.4 mM isopropyl β-D-1-thiogalactopyranoside for 20 hrs at 20°C. Cells were harvested by centrifugation and resuspended in binding buffer [25 mM Tris·HCl pH = 8.0, 150 mM NaCl, 5 mM β-mercaptoethanol, 20 mM imidazole, and Xpert protease inhibitor mixture (GenDEPOT)]. Cells were disrupted by sonication and cell debris were removed by centrifugation. The supernatant was loaded onto a Ni2+ Sepharose 6 Fast Flow column. After washing, protein was eluted by adding buffer A (25 mM Tris·HCl, pH 8.0, 150 mM NaCl, 5 mM β-mercaptoethanol, and Xpert protease inhibitor mixture) supplemented with 40, 60, 80, and 100 mM imidazole, respectively. Fractions containing M^pro^ based on SDS-PAGE were pooled and buffer-exchanged with buffer B [20 mM Tris·HCl, pH 7.3, 150 mM NaCl, 1 mM EDTA and 1 mM dithiothreitol (DTT)] using a Amicon Ultra-15 Centrifugal Filter Unit (Millipore Sigma). The sample was treated with Precission protease to remove the His-tag and create an authentic M^pro^ C-terminus. The sample was then applied to a Ni^2+^ Sepharose 6 Fast Flow resin was applied to the protein solution to remove remaining M^pro^-His. M^pro^ was then concentrated and applied to a Superdex 75 Increase 10/300GL size-exclusion column (GE Healthcare) preequilibrated with buffer B. Fractions containing M^pro^ were assessed by SDS-PAGE and pooled for further use.

### Enzyme kinetics and inhibition assays *in vitro*

To determine Michaelis-Menten kinetic parameters we utilized a fluorescent peptide, Dabcyl-KTSAVLQSGFRKM-E(Edans)-NH2, as a FRET substrate with an excitation wavelength of 360 nm and emission wavelength of 460 nm as described (*22*). We first determined the inner filter effect associated with the peptide and hydrolysis. This was done by measuring the fluorescence of the FRET peptide at 0, 7.5, 15, 30, 60, 90, and 120 µM and also at these peptide concentrations in the presence of 0.5 µM free EDANS. The difference in fluorescence with and without EDANS was used to calculate an inner filter effect correction as described (*40*). Fluorescence units were converted to hydrolysis product concentration in µM by running the M^pro^ hydrolysis reaction to completion for 0.5, 1, 5, 10, and 15 µM of FRET peptide and determining the change in fluorescence for the reaction, after inner filter effect correction. The change in fluorescence for the complete reaction was plotted versus peptide concentration to create a standard curve to convert fluorescence units to mM peptide hydrolyzed.

The Michaelis-Menten kinetic parameters *k*_cat_, *K*_M_ and *k*_cat_/*K*_M_ were determined for M^pro^ and mutant derivatives by measuring the initial velocity of peptide hydrolysis as indicated by the change in fluorescence and converted to µM. An inner filter effect correction was determined for each peptide concentration used and was applied to determine the initial velocity. The reactions were performed in 20 mM Tris-Cl pH 7.3, 100 mM NaCl, 1 mM EDTA, 1 mM DTT, and 0.02% Tween-20. The initial velocities were plotted versus peptide concentration and fit to the Michaelis-Menten equation:

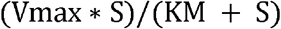

with *k*_cat_ = V_max_/E_0_, where E_0_ is the enzyme concentration used.

The potency of the PF07321332 and S217622 compounds for inhibition of M^pro^ and the A173V and ΔP168/A173V enzymes was evaluated using hydrolysis of the Dabcyl-KTSAVLQSGFRKM-E(Edans)-NH2 FRET peptide described above as a reporter. The inhibition assays were performed in 20 mM Tris-Cl pH 7.3, 100 mM NaCl, 1 mM EDTA, 1 mM DTT, and 0.02% Tween-20. The reporter peptide was used at 15 µM with increasing concentrations of compound to evaluate inhibition. Initial velocities of peptide hydrolysis at increasing inhibitor concentrations were performed in triplicate and the resulting average and standard deviation was fit to the Morrison equation for tight binding inhibitors (*41*) to obtain a *K*_i_ value. The error on the *K*_i_ value is determined from standard error of fitting to the equation. The K_M_ value used for the Morrison equation was from the Michaelis-Menten kinetic analysis with the FRET peptide for M^pro^ and each mutant as determined in this study.

Morrison equation:

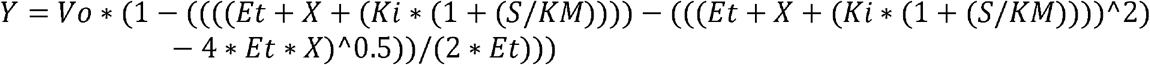

where Y is enzyme activity, X is the concentration of inhibitor, Et is the concentration of enzyme, K_m_ is the Michaelis constant, and S is the substrate concentration.

### Production of recombinant SARS-CoV-2

Recombinant SARS2 containing the M^pro^ variants of interested were generated using a previously described bacterial artificial chromosome (BAC)-based reverse genetics system based on the USA-WA1/2020 (WA-1) strain (accession no. MN985325), referred to as rSARS2/mCherry (*25, 42*). To introduce the desired mutation into full length SARS-CoV-2, first a plasmid containing the ORF1a which encodes Nsp5 (termed pUC57-F3) was used as a template for site directed mutagenesis to introduce the ΔP168, A173V and ΔP168/A173V mutations which were confirmed by Sanger sequencing. Next, the fragment containing the mutations of interested were inserted into the full length rSARS2 BAC by using the MluI and PacI restriction enzymes. Vero E6 cells expressing hACE2 and TMPRSS2 (Vero AT) were obtained from BEI Resources (NR-54970) and maintained in DMEM with 10% FBS, 1% PSG, and 10 µg/ml puromycin. For virus rescue, the full-length BAC containing the M^pro^ mutants were transfected into confluent monolayers of Vero AT cells (10^6^ cells/well, 6-well plate format). At 24hpi, the media was changed to post-infection medium (DMEM supplemented with 2% FBS and 1% PSG) and at 48hpi the cells were split into a T75 flask. After an additional 72h incubation, cell culture supernatants were harvested, labeled as P0 and frozen at −80° C. Monolayers of Vero AT cells (indicate the number of cells and format) were infected at low multiplicity of infection (MOI 0.0001) with P0 for 48h to generate P1 stocks. Viral RNA was extracted from P1 viral stocks and subjected to Illumina next generation sequencing (NGS) to confirm the presence of the desired mutations within Nsp5. P1 virus stocks were titrated and used for downstream antiviral and growth kinetic assays.

### Antiviral and growth kinetic assays

To avoid the nirmatrelvir efflux seen in Vero E6 cells, human A549-hACE2 cells (5×10^4^ cells/well, 96-well plate format, quadruplicates) were infected with 200 PFU/well of WT, ΔP168, A173V, or ΔP168/A173V rSARS2/mCherry and incubated for 1h at 37°C in a 5% CO_2_ incubator. After virus adsorption, cells were washed with PBS and incubated at 37°C in phenol red-free post-infection media containing two-fold serial dilutions of the indicated antiviral drug (nirmatrelvir, ensitrelvir, or remdesivir with a starting concentration of 25 µM). Fluorescence mCherry expression was determined at 72 hpi using a fluorescence microscope (EVOS M5000) and a Synergy LX Multimode plate reader (Agilent). Fluorescence values of mCherry virus-infected cells in the absence of antiviral were used to calculate 100% viral infection. Cells in the absence of viral infection were used to calculate the fluorescence background. The 50% inhibitory concentrations (IC_50_) were determined with a sigmoidal dose response curve (Graphpad Prism 9).

To determine viral growth kinetics of the WT and mutant rSARS-CoV-2, monolayers of Vero E6 cells (4×10^5^ cells/well, 12-well plate format, triplicates) were infected (MOI 0.01) and incubated for 1 hr at 37°C in a 5% CO_2_ incubator. After virus adsorption, cells were washed with PBS and incubated at 37°C in post-infection media At each of the indicated time points (12, 24, 48, and 72 hpi), viral titers were determined in cell culture supernatants by standard plaque assay.

### Structural modeling

#### Model construction

##### WT

Washington strain (*i*.*e*., “original” 2019 strain; wildtype, WT) M^pro^ structure was prepared for simulation from Protein Data Bank deposition 7BB2(*43*). In chains A and B of 7BB2, Cys128 is resolved in two conformations, thus care was taken to remove the B conformation of Cys128 from both chains by manual manipulation of the .pdb file and then renumbering atoms with pdb-tools (*44*). PROPKA3 was then used to calculate protonation states of titratable residues in the WT structure (*45*). At pH 7.4, deprotonated protonation states for all aspartate and glutamate were deemed appropriate, as well as protonated states for all lysines, tyrosines, arginines, and cysteines not involved in disulfide bonding. No cysteines were resolved in disulfide bonding patterns (*43*). Using atomic positions from 7BB2 structure deposited within the protein data bank, histidine residues 80 and 164 were determined to be protonated at the Nδ atom, whereas histidine residues 64, 163, 172, and 246 were determined to be protonated at the Nε atom. However, upon manual investigation of M^pro^’s mechanism as well as residue arrangement in the binding cleft, we determined the protonation state of histidine 41 needed to be switched from protonation on the Nε atom (as deposited in the PDB) to protonation on the Nδ atom. Per residue protonation state decisions are listed in **Table S1**. Molefacture, a tool within Visual Molecular Dynamics (VMD), was then used to rotate chain A Cys145 and chain A His 41 such that His41’s deprotonated Nε was pointed to Cys145’s sulfhydryl group, *i*.*e*., a conformation suitable for M^pro^ catalytic function (*46*). This molefacture procedure was repeated for chain B, to ensure both catalytic dyads began simulation in catalytically active conformation. The structure was then parsed with VMD checking tools to ensure there were no chirality errors or cis-peptide bonds. VMD tool psfgen was then used to construct a protein structure file (psf) from the resultant pdb file. This final structure was then also used as the base structure for building the A173V M^pro^ variant. Due to the fact that chain A of PDB ID 7SI9 was resolved with 305 residues (residues 1 to 306) but chain B was resolved with 299 residues (residues 1 to 300), we constructed M^pro^ in this asymmetric way as well. All variants constructed herein were identically asymmetrical, *i*.*e*., all chain A’s contain 305 residues and all chain B’s contain 299 residues.

##### A173V

Starting from the constructed WT pdb file, VMD tool psfgen was used along with the mutate command to change A173 to V173.

##### ΔP168

RoseTTAFold (*47, 48*) was used to construct a homology model of M^pro^ ΔP168 with nirmatrelvir-bound structure PDB ID 7SI9 (*19*) as the template. PROPKA3.0 was again used to calculate pKa’s of all titratable residues in M^pro^. The same protonation scheme was deemed appropriate for all residues (*i*.*e*., deprotonation of all aspartates/glutamates, protonation of all lysines/tyrosines/cysteines/arginines) and the same histidine protonation scheme was adopted as resolved from 7BB2 (with exception of His41 which was set to Nδ protonation). Like in the WT model construction, molefacture was used to rotate Cys145 and His41 such that His41’s deprotonated Nε was pointed to Cys145’s sulfhydryl group, *i*.*e*., a conformation suitable for M^pro^ catalytic function (procedure done for dyads on both chains A and B). The structure was then parsed with VMD checking tools to ensure there were no chirality errors or cis-peptide bonds. VMD tool psfgen was then used to construct a protein structure file (psf) from the resultant pdb. The resultant structure was also used as the base structure for building the ΔP168/A173V M^pro^ variant.

##### ΔP168/A173V

Starting from the constructed ΔP168 M^pro^ variant structure, VMD tool psfgen was used along with the mutate command to add A173V.

##### Solvation and Neutralization

VMD was used to solvate all M^pro^ variant structures in water boxes of 90.7 × 97.2 × 114.3 Å^3^ size and neutralized with 150 mM NaCl. Total number of atoms and exact system box sizes per variant can be seen in **Table S2**.

### Molecular Dynamics Simulations

For all MD simulations described herein, the following force field parameters were used. All atoms in all systems were described according to the CHARMM36m force field (*49, 50*). All water molecules were described according to the TIP3P water model (*51*). Infinite bulk conditions were modeled with periodic boundary conditions (PBC) and long-range electrostatic interactions were calculated with Particle Mesh Ewald (PME, interpolation order 8, grid spacing 2.0). Nonbonded atom pair lists were generated for all atoms within 15.5 Å of one another. Nonbonded interaction energies were calculated for all atoms within 10 Å of one another. For nonbonded atom pairs beyond 12Å from one another, nonbonded interactions were assigned a zero-energy contribution and not calculated. For atom pairs between 10 and 12Å from one another, a switching function was used to gradually switch nonbonded terms from their calculated value to zero energetic contribution. The SHAKE algorithm was applied to constrain all bonds between heavy atoms and hydrogens to their equilibrium distance value as listed in CHARMM36m parameter files. 1-2 and 1-3 nonbonded interactions were excluded (not calculated), while electrostatic interactions of 1-4 pairs are scaled (default scaling factor 1.0) and Lennard-Jones potentials were modified according to CHARMM36m parameter files. Unless otherwise noted, NAMD2.14 was used to conduct all following MD simulations (*52, 53*). For a summary of all MD simulation steps described below, see **Table S3**. All MD simulations were performed with San Diego Supercomputing Network’s Hopper GPU cluster.

#### Minimization

While holding catalytic dyad residues (Cys145 and His41) in position with a constraint (*i*.*e*., “fixed atoms”), we launched three replicas of 10,000 steps of conjugate gradient and line search algorithm minimization for each M^pro^ variant. We then followed each of these minimization procedures with 500 steps of dynamics at 310 K. All subsequent MD simulation steps were launched from final coordinates and velocities of these 500 steps of 310 K dynamics, and thus all following simulation steps were performed in triplicate.

#### Heating

Velocities from previous 500 steps at 310 K were used to launch to heating procedure in which the Langevin temperature and piston temperature were gradually increased from 10K to 310 K in increments of 25 K with 10,080 MD steps performed at each temperature (at 1fs/timestep, thus 10ps of sampling per temperature). Once the system reached 310 K, an additional 10,080 steps (10ps) of simulation were performed (thus 20 ps total performed at 310 K). During this heating procedure, catalytic dyad residues were again constrained (*i*.*e*., Cys145 and His41 of chains A and B were held fixed) to their final position after the previous minimization step.

#### NpT Equilibration

Final coordinates and velocities from heating simulations were used to launch 25,200 steps (1fs/timestep, thus 252 ps) of NpT equilibration. Pressure was set to 1 atmosphere, and the periodic cell was allowed to be flexible during simulation (useFlexibleCell set to yes) to equilibrate the cell volume. During NpT equilibration, constraint on the catalytic dyad residues were removed and instead the catalytic dyad residues were restrained to their positions following heating (harmonic restraint with force constant of 1 kcal/mol/Å).

#### NVT Equilibration

Final coordinates and velocities from NpT equilibration simulations were used to launch 27,500,000 steps (1 fs/timestep, thus 27.5 ns) of NVT equilibration. During NVT equilibration, pressure was maintained at 1 atmosphere. The box dimensions were fixed to the dimensions from the final step of NpT equilibration (useFlexibleCell set to no). All restraints and constraints were removed, and no new constraints/restraints were implemented during NVT equilibration, thus all atoms were allowed full flexibility.

#### Statistical Sampling with NAMD3.0/GPU

12 Final coordinates and velocities from NVT equilibration simulations were used to launch 275,000,000 steps (2 fs/timestep, thus 550 ns; note the timestep change from previous methods) of statistically relevant sampling at the NVT ensemble. Pressure was maintained at 1 atmosphere, and the volume was fixed at each step (useFlexibleCell set to no). Due to the switch from 1 to 2 fs timestep, and the switch from NAMD2.14 on CPUS to NAMD3.0 on GPU, we simulated for 550 ns to allow for 50 ns of final “equilibration” if need be, before collecting results from the final 500 ns. However, after performing initial analysis, we observed that all simulations appeared similarly equilibrated by Root Mean Square Deviation from their starting structure (**Figure S6**), thus we deemed it appropriate to incorporate all 550ns sampling per replica per M^pro^ variant. Every 2500th frame was written to a dcd file for analysis, thus from 275,000,000 steps we collected 110,000 frames for analysis.

### Computational analyses

MDAnalysis tools were used for all analyses described below (*54, 55*). All M^pro^ trajectories were stripped of water and ion atoms. Due to periodic boundary effects, chains A and B had to be split and analyzed separately. Each M^pro^ trajectory was then aligned to its first frame by Cα atoms to cancel rotational and translational degrees of freedom.

#### Root mean square deviation (RMSD)

Root mean square deviation calculations were performed with the MDAnalysis RMSD calculator. First, each trajectory was realigned (for each variant, each chain, and each replica) according to the positions of residues 1 to 167 and 169 to 300 to their position in the first frame, to ensure no biases introduced by the addition of the extra residue 168 in the alignment. Then RMSD was calculated for each of these same Cα atoms and is plotted in **Figure S6**. To monitor trends more clearly in data, we then calculated a rolling average of the RMSD over the whole data set. In **Figure S6** we have plotted both the full data set, in transparent blue and brown lines, and the rolling average of the data set, in dark blue and brown lines. To calculate the average RMSD trend, for each timestep we calculated the average RMSD at that timepoint across all replicas and across both chains. For the average RMSDs, we have again plotted the full data set, in transparent black lines, and the rolling average of average RMSDs, in dark black lines.

#### Root mean square fluctuation (RMSF)

To calculate root mean square fluctuations (RMSFs) we first aligned each trajectory (per variant, per replica, and residues 1-300 per chain chain) according to Ca atomic positions in the starting frame. We then used the MDAnalysis RMSF tool to calculate the per residue fluctuation over the course of each trajectory. For each residue, we then averaged the RMSF of that residue and used the standard deviation in RMSFs per residue to estimate error. To ensure that all residues were aligned when plotting RMSFs per residue, in variants where residue 168 was deleted we interpolated from average RMSFs of residue 167 and 169.

#### Hydrogen bonding analysis

To calculate percentage of frames with a “hydrogen bond” between backbone atoms of L167 and G170, for each chain and each replica, we counted all the frames for which the following two requirements were satisfied: (1) the distance between L167 backbone oxygen and G170 backbone nitrogen were within 4Å of one another and (2) the angle formed between the L167 backbone oxygen, G170 backbone hydrogen, and G170 backbone nitrogen was greater than 120º. We then determined the percentage of frames that exhibited a hydrogen bond at this position and averaged these percentages over all replicas and across the two chains. We then used the standard deviations of these averages to estimate error in our percentage calculations.

#### S4 subsite penetration by M49

To calculate the degree of S4 subsite penetration by residue M49, for every frame in all simulations (*i*.*e*., each variant, each replica, for both chains) we calculated the distance between the center of mass of residue M49 and the center of mass of residues 165-167 and 188-192. We then used the gaussian_kde (kernel density estimation with Gaussian kernels) module in SciPy to estimate smooth density plots from histograms of these M49-S4 subsite distances.

### SARS-CoV-2 variant analyses

Relative distributions of amino acid changes at the amino acid positions of interest in M^pro^ were counted using the GISAID EpiCoV web server and filtered based on viral genome sequences that are considered “Complete” and “High Coverage”. The frequency of each amino individual amino acid and in frame deletions were divided by the total number of changes found at said position to calculate the relative frequency and plotted as a heatmap using Graphpad Prism 9. To generate phylogenetic trees of viral genomes containing the M^pro^ variants of interest, full length viral genomes were first retrieved from the GISAID database and filtered to exclude sequences with low coverage and sequences that contained obvious errors or poor coverage within Nsp5 were manually removed. Viral genomes were then uploaded to the UShER and placed within a phylogenetic tree of all available sequences in the GISAID database with the number of samples per subtree showing sample placement set to 50. The generated phylogenetic trees were visualized using Auspice.us from NextStrain (*56*). Metadata for the viral genomes were retrieved from GISAID and overlayed on the phylogenetic tree using the Auspice.us web application. Viral genomes were similarly filtered to determine the overall frequency of each variant with additional filtering of separating all Omicron sequences compared to all previous lineages.

## List of Supplementary Materials

Fig S1 to S12

Table S1 to S3

## Acknowledgments

We thank several members of the Harris lab and R. Langlois for support and constructive feedback. RSH is an Investigator of the Howard Hughes Medical Institute, a CPRIT Scholar, and the Ewing Halsell President’s Council Distinguished Chair at University of Texas Health San Antonio.

## Author contributions

Conceptualization: SAM, RSH

Methodology: SAM, EH, AK, CN, FLK, CY, SNM, TP

Investigation: SAM, EH, AK, CN, FLK, CY, SNM, FLK, FC, MAE

Visualization: SAM, FLK, HA

Funding acquisition: SAM, EH, DvL, RSH, REA, TP, LMS

Project administration: RSH

Supervision: RSH, DvL, REA, TP, LMS

Writing – original draft: SAM, RSH

Writing – review & editing: All authors

## Data and materials availability

All data are available in the main text or the supplementary materials.

## References and Notes

1. K. Anand, J. Ziebuhr, P. Wadhwani, J. R. Mesters, R. Hilgenfeld, Coronavirus main proteinase (3CLpro) structure: basis for design of anti-SARS drugs. Science 300, 1763–1767 (2003).

2. L. Zhang et al., Crystal structure of SARS-CoV-2 main protease provides a basis for design of improved alpha-ketoamide inhibitors. Science 368, 409–412 (2020).

3. B. Boras et al., Preclinical characterization of an intravenous coronavirus 3CL protease inhibitor for the potential treatment of COVID19. Nat Commun 12, 6055 (2021).

4. R. F. Service, A call to arms. Science 371, 1092–1095 (2021).

5. C. Flexner, HIV-protease inhibitors. N Engl J Med 338, 1281–1292 (1998).

6. J. Anderson, C. Schiffer, S. K. Lee, R. Swanstrom, Viral protease inhibitors. Handbook of Experimental Pharmacology, 85–110 (2009).

7. A. Luttens et al., Ultralarge virtual screening identifies SARS-CoV-2 main protease inhibitors with broad-spectrum activity against coronaviruses. J Am Chem Soc 144, 2905–2920 (2022).

8. C. H. Zhang et al., Potent noncovalent inhibitors of the main protease of SARS-CoV-2 from molecular sculpting of the drug perampanel guided by free energy perturbation calculations. ACS Cent Sci 7, 467–475 (2021).

9. J. Qiao et al., SARS-CoV-2 M(pro) inhibitors with antiviral activity in a transgenic mouse model. Science 371, 1374–1378 (2021).

10. A. D. Rathnayake et al., 3C-like protease inhibitors block coronavirus replication in vitro and improve survival in MERS-CoV-infected mice. Sci Transl Med 12, (2020).

11. C. M. Consortium, Open science discovery of oral non-covalent SARS-CoV-2 main protease inhibitor therapeutics. BioRxiv https://doi.org/10.1101/2020.10.29.339317 (2022).

12. D. R. Owen et al., An oral SARS-CoV-2 M(pro) inhibitor clinical candidate for the treatment of COVID-19. Science 374, 1586–1593 (2021).

13. https://www.shionogi.com/global/en/news/2022/11/e20221122.html

14. Y. Unoh et al., Discovery of S-217622, a noncovalent oral SARS-CoV-2 3CL protease inhibitor clinical candidate for treating COVID-19. Journal of Medicinal Chemistry 65, 6499–6512 (2022).

15. A. Telenti, E. B. Hodcroft, D. L. Robertson, The evolution and biology of SARS-CoV-2 variants. Cold Spring Harbor Perspectives in Medicine 12, (2022).

16. S. A. Moghadasi et al., Gain-of-signal assays for probing inhibition of SARS-CoV-2 M(pro)/3CL(pro) in living cells. mBio 13, e0078422 (2022).

17. https://www.fda.gov/media/155050/download

18. Y. Shu, J. McCauley, GISAID: Global initiative on sharing all influenza data - from vision to reality. Euro Surveill 22, (2017).

19. D. W. Kneller et al., Covalent narlaprevir- and boceprevir-derived hybrid inhibitors of SARS-CoV-2 main protease. Nat Commun 13, 2268 (2022).

20. J. Barrila, U. Bacha, E. Freire, Long-range cooperative interactions modulate dimerization in SARS 3CLpro. Biochemistry 45, 14908–14916 (2006).

21. E. Heilmann et al., A VSV-based assay quantifies coronavirus Mpro/3CLpro/Nsp5 main protease activity and chemical inhibition. Commun Biol 5, 391 (2022).

22. S. Chamakuri et al., DNA-encoded chemistry technology yields expedient access to SARS-CoV-2 M(pro) inhibitors. Proc Natl Acad Sci U S A 118, (2021).

23. S. Legare, F. Heide, B. A. Bailey-Elkin, J. Stetefeld, Improved SARS-CoV-2 main protease high-throughput screening assay using a 5-carboxyfluorescein substrate. J Biol Chem 298, 101739 (2022).

24. C. J. Kuo, Y. H. Chi, J. T. Hsu, P. H. Liang, Characterization of SARS main protease and inhibitor assay using a fluorogenic substrate. Biochem Biophys Res Commun 318, 862–867 (2004).

25. K. Chiem et al., Generation and characterization of recombinant SARS-CoV-2 expressing reporter genes. J Virol 95, (2021).

26. Y. Turakhia et al., Ultrafast sample placement on existing tRees (UShER) enables real-time phylogenetics for the SARS-CoV-2 pandemic. Nat Genet 53, 809–816 (2021).

27. Z. Lv et al., Targeting SARS-CoV-2 proteases for COVID-19 antiviral development. Front Chem 9, 819165 (2021).

28. Y. Hu et al., Naturally occurring mutations of SARS-CoV-2 main protease confer drug resistance to nirmatrelvir. BioRxiv https://doi.org/10.1101/2022.06.28.497978 (2022).

29. Y. Zhou et al., Nirmatrelvir resistant SARS-CoV-2 variants with high fitness in vitro. BioRxiv https://doi.org/10.1101/2022.06.06.494921 (2022).

30. S. Iketani et al., Multiple pathways for SARS-CoV-2 resistance to nirmatrelvir. BioRxiv, (2022).

31. D. Jochmans et al., The substitutions L50F, E166A and L167F in SARS-CoV-2 3CLpro are selected by a protease inhibitor in vitro and confer resistance to nirmatrelvir. BioRxiv https://doi.org/10.1101/2022.06.07.495116 (2022).

32. E. Heilmann, F. Costacurta, A. Volland, D. von Laer, SARS-CoV-2 3CLpro mutations confer resistance to Paxlovid (nirmatrelvir/ritonavir) in a VSV-based, non-gain-of-function system. BioRxiv https://doi.org/10.1101/2022.07.02.495455 (2022).

33. Y. Zhao et al., Structural basis for replicase polyprotein cleavage and substrate specificity of main protease from SARS-CoV-2. Proc Natl Acad Sci U S A 119, e2117142119 (2022).

34. https://www.japic.or.jp/mail_s/pdf/23-11-1-07.pdf

35. G. Dias Noske et al., Structural basis of nirmatrelvir and ensitrelvir resistance profiles against SARS-CoV-2 main protease naturally occurring polymorphisms. BioRxiv https://doi.org/10.1101/2022.08.31.506107 (2022).

36. R. A. Bull et al., Analytical validity of nanopore sequencing for rapid SARS-CoV-2 genome analysis. Nat Commun 11, 6272 (2020).

37. E. Lontok et al., Hepatitis C virus drug resistance-associated substitutions: State of the art summary. Hepatology 62, 1623–1632 (2015).

38. N. M. King, M. Prabu-Jeyabalan, E. A. Nalivaika, C. A. Schiffer, Combating susceptibility to drug resistance: lessons from HIV-1 protease. Chem Biol 11, 1333–1338 (2004).

39. A. Molla et al., Ordered accumulation of mutations in HIV protease confers resistance to ritonavir. Nat Med 2, 760–766 (1996).

40. Y. Liu et al., Use of a fluorescence plate reader for measuring kinetic parameters with inner filter effect correction. Analytical Biochemistry 267, 331–335 (1999).

41. J. F. Morrison, Kinetics of the reversible inhibition of enzyme-catalysed reactions by tight-binding inhibitors. Biochim Biophys Acta 185, 269–286 (1969).

42. C. Ye, L. Martinez-Sobrido, Use of a bacterial artificial chromosome to generate recombinant SARS-CoV-2 expressing robust levels of reporter genes. Microbiol Spectr 10, e0273222 (2022).

43. E. Costanzi et al., Structural and biochemical analysis of the dual inhibition of MG-132 against SARS-CoV-2 main protease (Mpro/3CLpro) and human cathepsin-L. International journal of Molecular Sciences 22, (2021).

44. J. Rodrigues, J. M. C. Teixeira, M. Trellet, A. Bonvin, pdb-tools: a swiss army knife for molecular structures. F1000Res 7, 1961 (2018).

45. M. H. Olsson, C. R. Sondergaard, M. Rostkowski, J. H. Jensen, PROPKA3: Consistent Treatment of Internal and Surface Residues in Empirical pKa Predictions. J Chem Theory Comput 7, 525–537 (2011).

46. W. Humphrey, A. Dalke, K. Schulten, VMD: visual molecular dynamics. Journal of Molecular Graphics 14, 33-38, 27-38 (1996).

47. I. R. Humphreys et al., Computed structures of core eukaryotic protein complexes. Science 374, eabm4805 (2021).

48. M. Baek et al., Accurate prediction of protein structures and interactions using a three-track neural network. Science 373, 871–876 (2021).

49. J. Huang, A. D. MacKerell, Jr., CHARMM36 all-atom additive protein force field: validation based on comparison to NMR data. Journal of Computational Chemistry 34, 2135–2145 (2013).

50. J. Huang et al., CHARMM36m: an improved force field for folded and intrinsically disordered proteins. Nat Methods 14, 71–73 (2017).

51. W. L. Jorgensen, J. Chandrasekhar, J. D. Madura, Comparison of simple potential functions for simulating liquid water. J Chem Phys 79, 926 (1983).

52. J. C. Phillips et al., Scalable molecular dynamics with NAMD. Journal of Computational Chemistry 26, 1781–1802 (2005).

53. J. C. Phillips et al., Scalable molecular dynamics on CPU and GPU architectures with NAMD. J Chem Phys 153, 044130 (2020).

54. N. Michaud-Agrawal, E. J. Denning, T. B. Woolf, O. Beckstein, MDAnalysis: a toolkit for the analysis of molecular dynamics simulations. Journal of Computational Chemistry 32, 2319–2327 (2011).

55. R. J. Gowers et al., MDAnalysis: a python package for the rapid analysis of molecular dynamics simulations. doi:10.25080/Majora-629e541a-00e (2019).

56. J. Hadfield et al., Nextstrain: real-time tracking of pathogen evolution. Bioinformatics 34, 4121–4123 (2018).

