## Supplementary material for "Transmissible SARS-CoV-2 variants with resistance to clinical protease inhibitors": Figs S1-S12 and Tables S1-S3

[*] Equal secondary contributions

**Contents:** Supplementary Figures S1-S12, Supplementary Tables S1-3

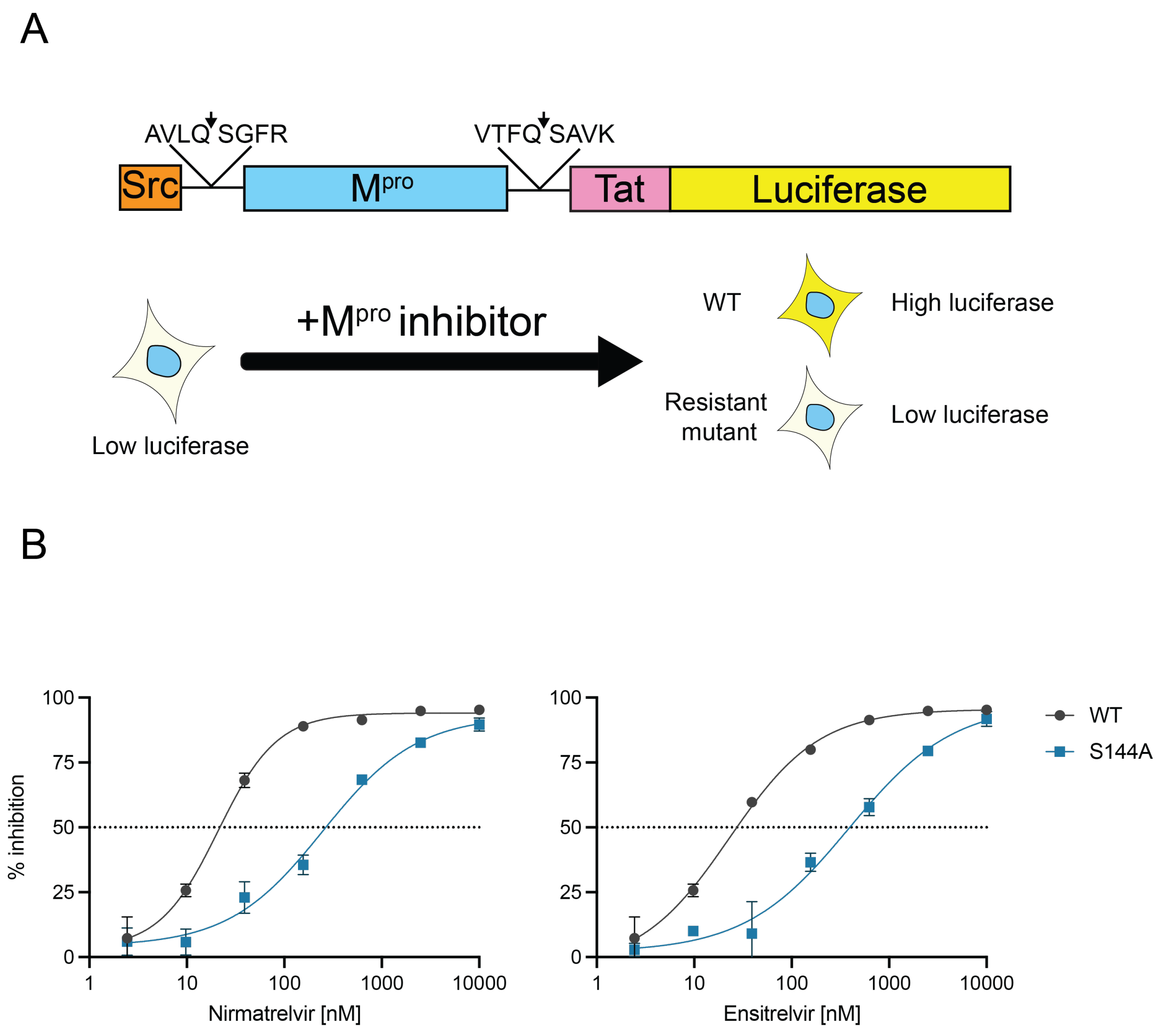


**Supplementary Figure 1. Live cell gain-of-signal assay for quantifying SARS2 $M^{pro}$ function.**

(**A**) Schematic of Src-$M^{pro}$-Tat-fLuc assay in which transfection of a catalytically active WT $M^{pro}$ construct into 293T cells yields low luciferase expression due to cleavage of host substrates that prevent reporter expression. Inhibition of $M^{pro}$ catalytic activity by chemical (shown) or genetic methods results in quantifiable increases in luminescent signal.

(**B**) Nirmatrelvir and ensitrelvir dose responses for WT and S144A $M^{pro}$ using the live cell Src-$M^{pro}$-Tat-fLuc assay (4-fold dilution series beginning at 10 µM; data are mean +/- SD of biologically independent triplicate experiments).

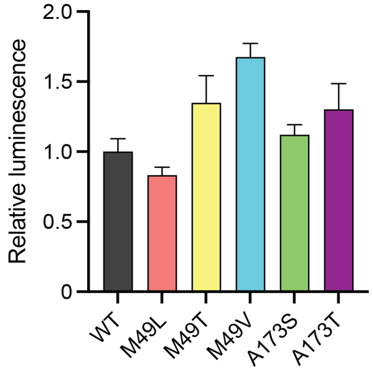


**Supplementary Figure 2. Background luminescence of M$^{pro}$ M49 and A173 variants.** Relative luminescence of 293T cells transfected with the Src-M$^{pro}$-Tat-fLuc reporter plasmid in the absence of inhibitor normalized to WT (data are mean +/- SD of 3 biologically independent triplicate experiments).

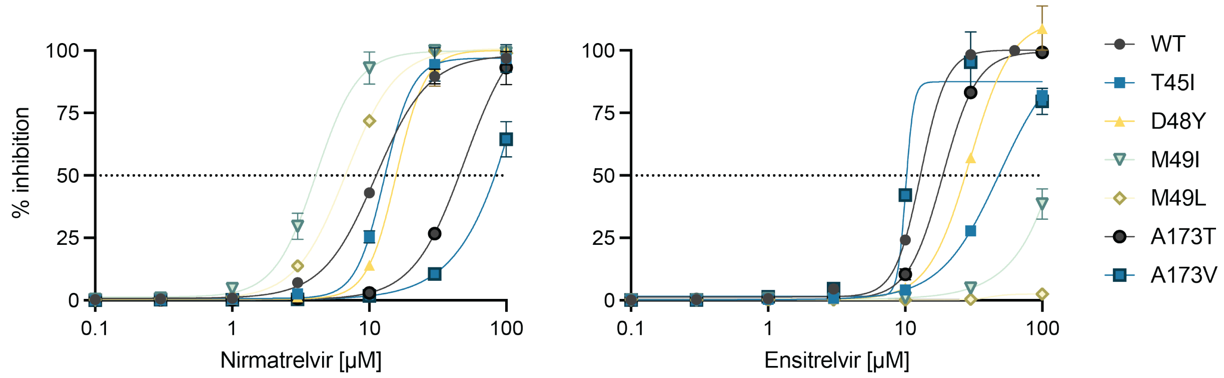


**Supplementary Figure 3. Orthologous validation of M^pro single mutant resistance phenotypes.**

Tests of the indicated M^pro single mutants against nirmatrelvir and ensitrelvir in a VSV-based M^pro *cis*-cleavage assay (3-fold dilution series starting at 100 µM; data are mean +/- SD of biologically independent triplicate experiments).

A

| $M^{pro}$ | $k_{cat}$ | $K_M$ | $k_{cat}/K_M$ |
|---|---|---|---|
| WT | 0.43 + 0.05 $s^{-1}$ | 57 + 15 µM | 7.5 x $10^3$ $M^{-1}s^{-1}$ |
| A173V | 1.2 + 0.25 $s^{-1}$ | 250 + 72 µM | 4.8 x $10^3$ $M^{-1}s^{-1}$ |
| ∆P168/A173V | 0.96 + 0.43 $s^{-1}$ | 190 + 130 µM | 5.1 x $10^3$ $M^{-1}s^{-1}$ |

B

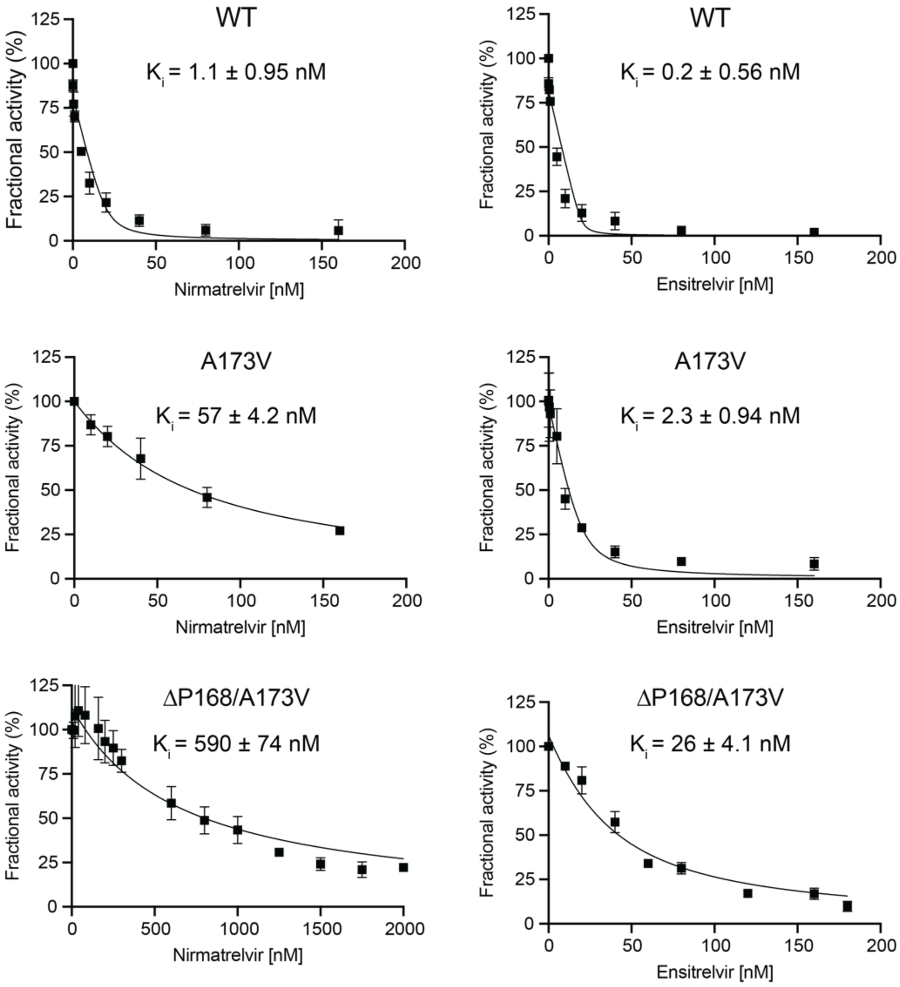


**Supplementary Figure 4**. **Activity and inhibition $M^{pro}$ A173V and ∆P168/A173V mutants using a FRET-based peptide cleavage assay *in vitro*.**

(**A**) Kinetic parameters of WT, A173V, and ∆P168/A173V mutants.

(**B**) Initial velocity for $M^{pro}$ hydrolysis of FRET peptide versus increasing nirmatrelvir and ensitrelvir concentrations. $K_i$ was determined by fit to the Morrison equation (Materials and Methods).

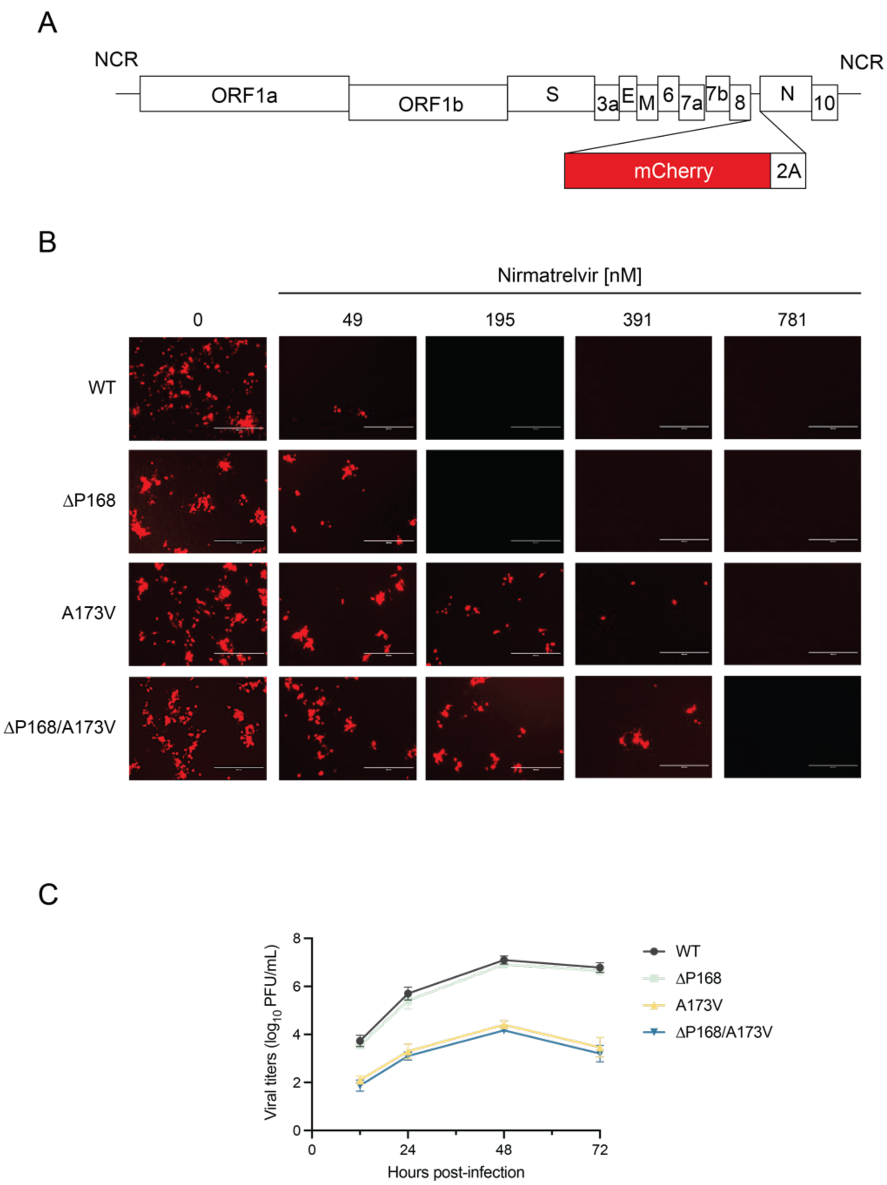


**Supplementary Figure 5**. **Recovery of mCherry expressing recombinant SARS-CoV-2 harboring M$^{pro}$ mutations.**

(**A**) Schematic of the BAC-based SARS-CoV-2 genome with a mCherry reporter separated from the nucleocapsid (*N*) by a 2A cleavage site.

(**B**) Representative fluorescent images of A549-hACE2 cells infected with recombinant SARS-CoV-2 harboring the indicated M$^{pro}$ variations in the presence of varying concentrations of nirmatrelvir (WT and P168/A173V images are identical to those in Fig. 3H and shown again here for comparison).

(**C**) Multicycle growth kinetics in in Vero-E6 cells of recombinant SARS-CoV-2 harboring the indicated M$^{pro}$ variations (mean values +/- SD from biological quadruplicate experiments).

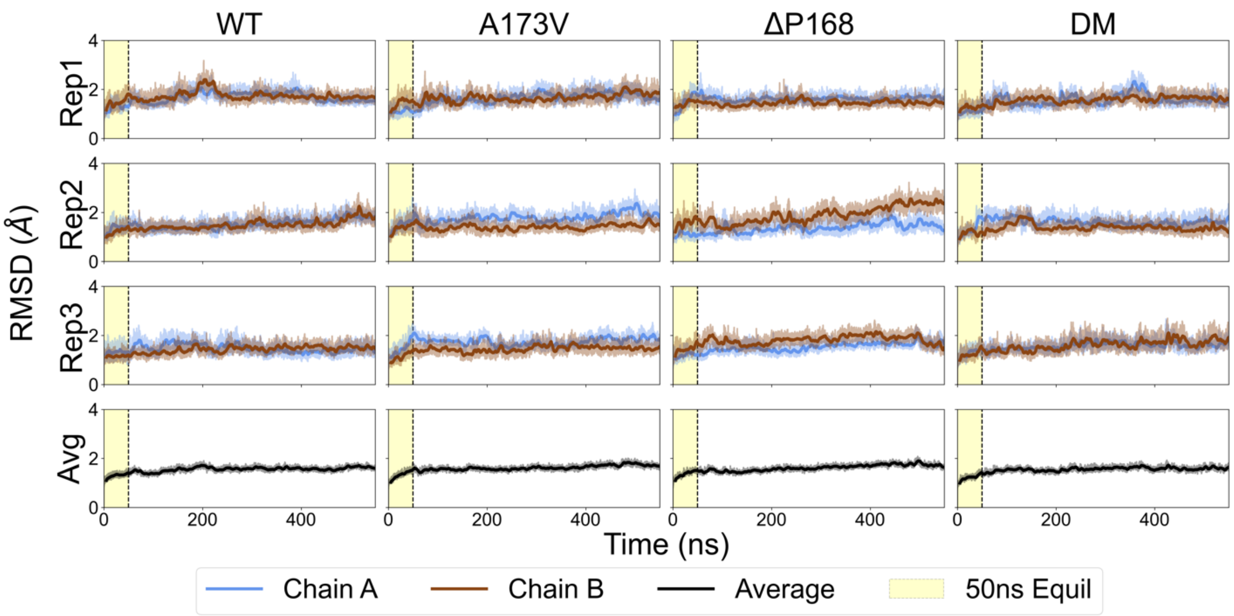


**Supplementary Figure 6. RMSD from MD simulations of M$^{pro}$ variants.**
RMSD plotted for all variants per replica, per chain, for all variants, as well as an average RMSD (bottom row of plots) which represents the average RMSD at each simulation step over all replicas and chains per variant. A yellow rectangle on each plot indicates the initial 50 ns simulation time we expected may be needed for equilibration, but RMSD data indicate these simulations are already sufficiently equilibrated (RMSDs all well below 4 Å) due to our extensive equilibration protocol already implemented (see Methods). Due to the fact that all RMSD values are low (<4Å) and plateaued (*i.e.*, no large drifts or trends over time), we can conclude that our M$^{pro}$ models are stably constructed and have been well equilibrated to temperature, pressure, and volume simulation parameters. Thus, trajectory frames collected from these 550 ns MD simulations can be reliably used to investigate M$^{pro}$ WT and variant structure/function/dynamics relationships.

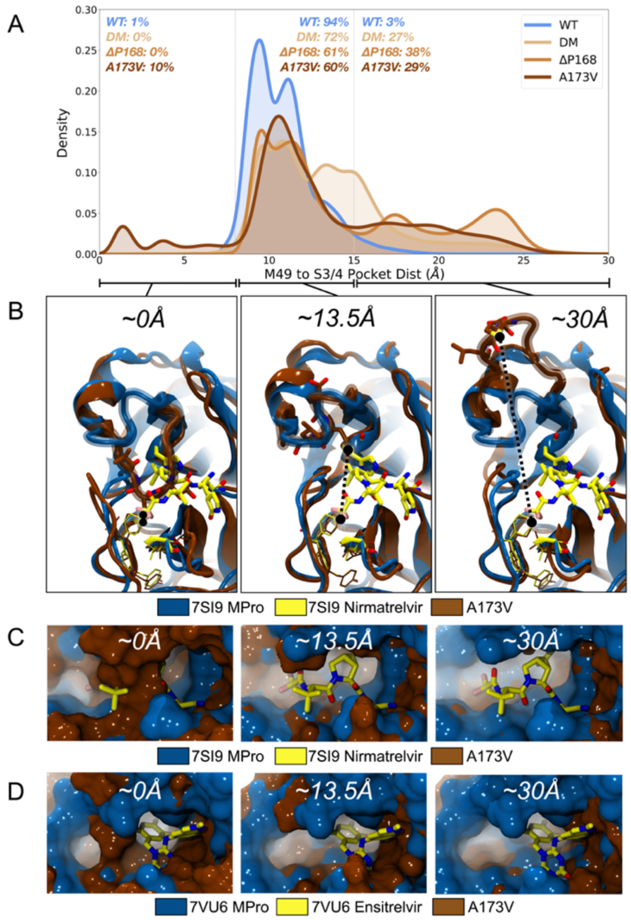


**Supplementary Figure 7: Mutation of A173V induces penetration of M49 into S4 pocket.**
(**A**) Plot demonstrating the distribution of frames in which M49 is found within the S3/S4 subpocket (<8Å from the S3/S4 subpocket), in a near crystal conformation (between 8 and 15Å from the S3/S4 subpocket), and wide open (beyond 15Å from the S3/S4 subpocket; plot same as main text Fig. 4A and shown again here for comparison to trajectory snapshots).
(**B**) Molecular models comparing snapshots from the A173V simulation (chocolate brown ribbons) to the 7SI9 crystal structure (blue ribbons) in which M49 is either penetrating the S3/S4 subpocket (labeled "~0Å"), in a near crystal conformation (labeled "~13.5A"), or wide open (labeled "~30Å").
(**C**) Molecular models comparing snapshots from the A173V simulation (chocolate brown surface) to the 7SI9 crystal structure (blue surface) with nirmatrelvir bound (carbon atoms shown in yellow licorice). When M49 penetrates the S3/S4 subpocket (labeled "~0Å") the nirmatrelvir binding mode is almost completely occupied, thus this conformation likely prevents nirmatrelvir binding. However, in the other two conformations (labeled "~13.5Å" and "~30Å"), the nirmatrelvir binding mode can be appropriately accommodated.
(**D**) Molecular models comparing snapshots from the A173V simulation (chocolate brown surface) to the 7VU6 crystal structure (blue surface) with ensitrelvir bound (carbon atoms shown in yellow licorice). When M49 penetrates the S3/S4 subpocket (labeled "~0Å") the ensitrelvir binding mode is still accommodated, likely allowing ensitrelvir binding. This difference in binding modes could explain why A173V and the double mutant variant are more resistant to nirmatrelvir than ensitrelvir.

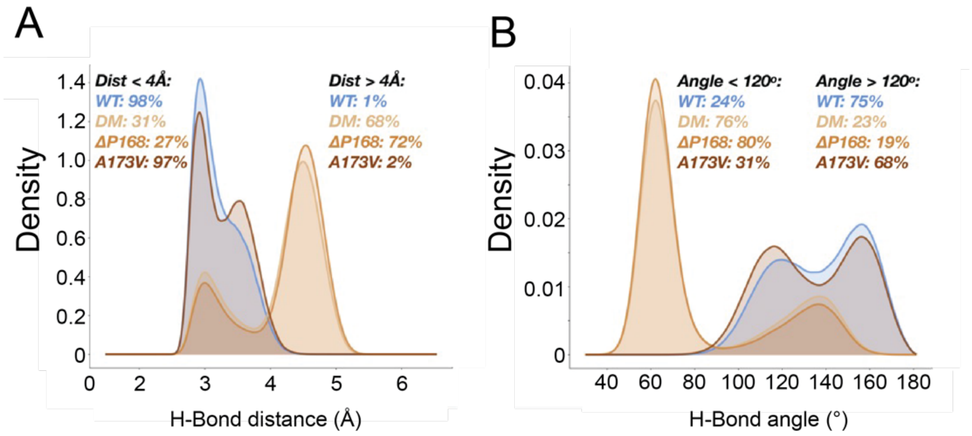


**Supplementary Figure 8. H-bonding pattern of L167-V171 backbone is perturbed by ΔP168.** (**A**) Distribution of distances and angles used to determine degree of hydrogen bonding between backbone atoms of residues L167 and G170. Distribution of distances between backbone carbonyl oxygen atom of L167 and backbone nitrogen atom of G170.

(**B**) Distribution of angles formed between the backbone carbonyl oxygen atom of L167, the backbone hydrogen atom of G170, and the backbone nitrogen atom of G170.

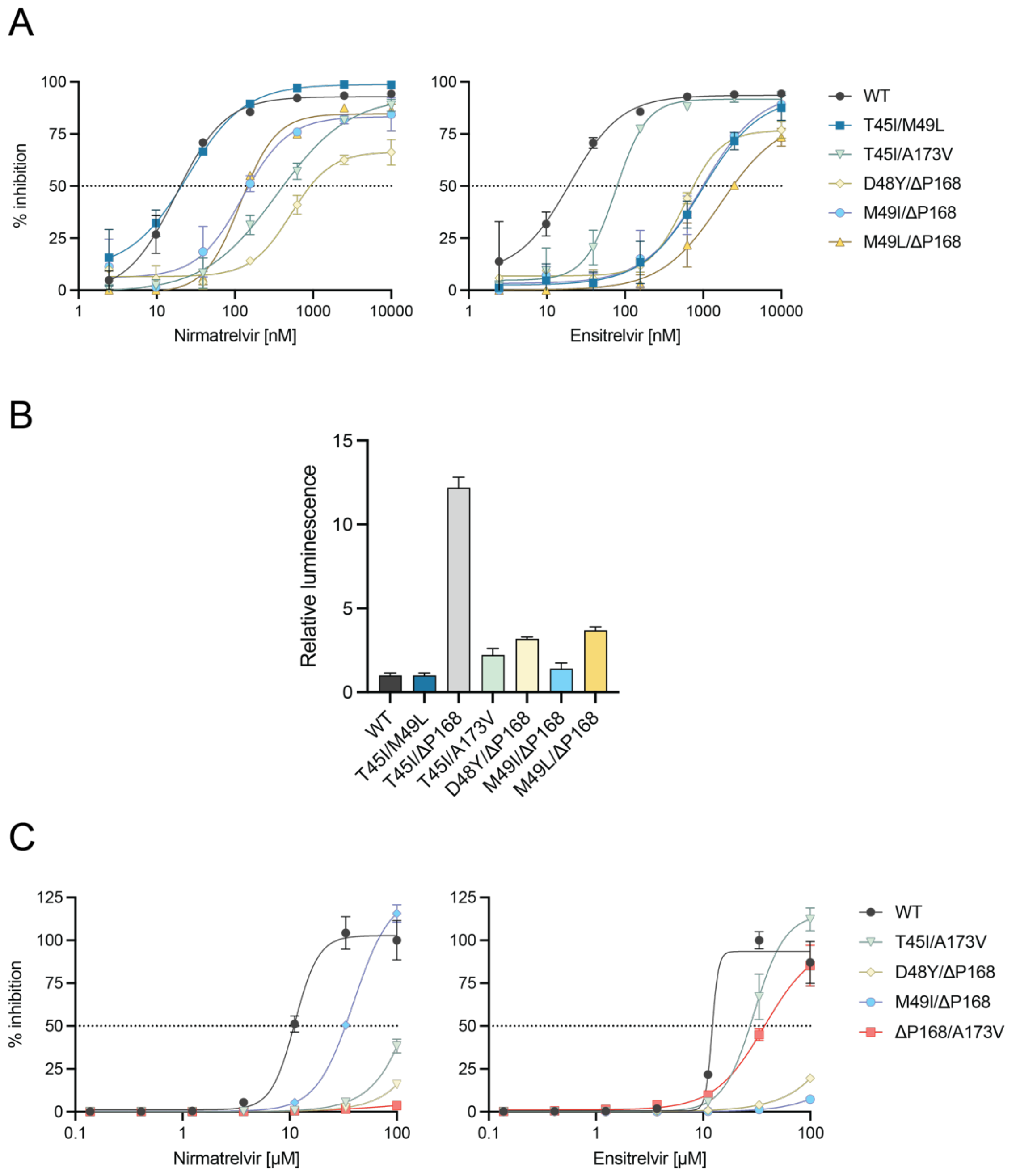


**Supplementary Figure 9. Analyses of additional double mutants using two live cell systems.** (**A**) Nirmatrelvir and ensitrelvir dose-responses for the indicated double mutants (4-fold dilutions beginning at 10 µM; data are mean +/- SD of biologically independent triplicate experiments).
(**B**) Background luminescence in the absence of drug relative to the WT as a proxy for catalytic activity (data are mean +/- SD of biologically independent triplicate experiments).
(**C**) Dose-responses of WT and double mutant $M^{pro}$ constructs for nirmatrelvir and ensitrelvir using a VSV-based $M^{pro}$ cis-cleavage assay (3-fold dilution series starting at 100 µM; data are mean +/- SD of biologically independent triplicate experiments).

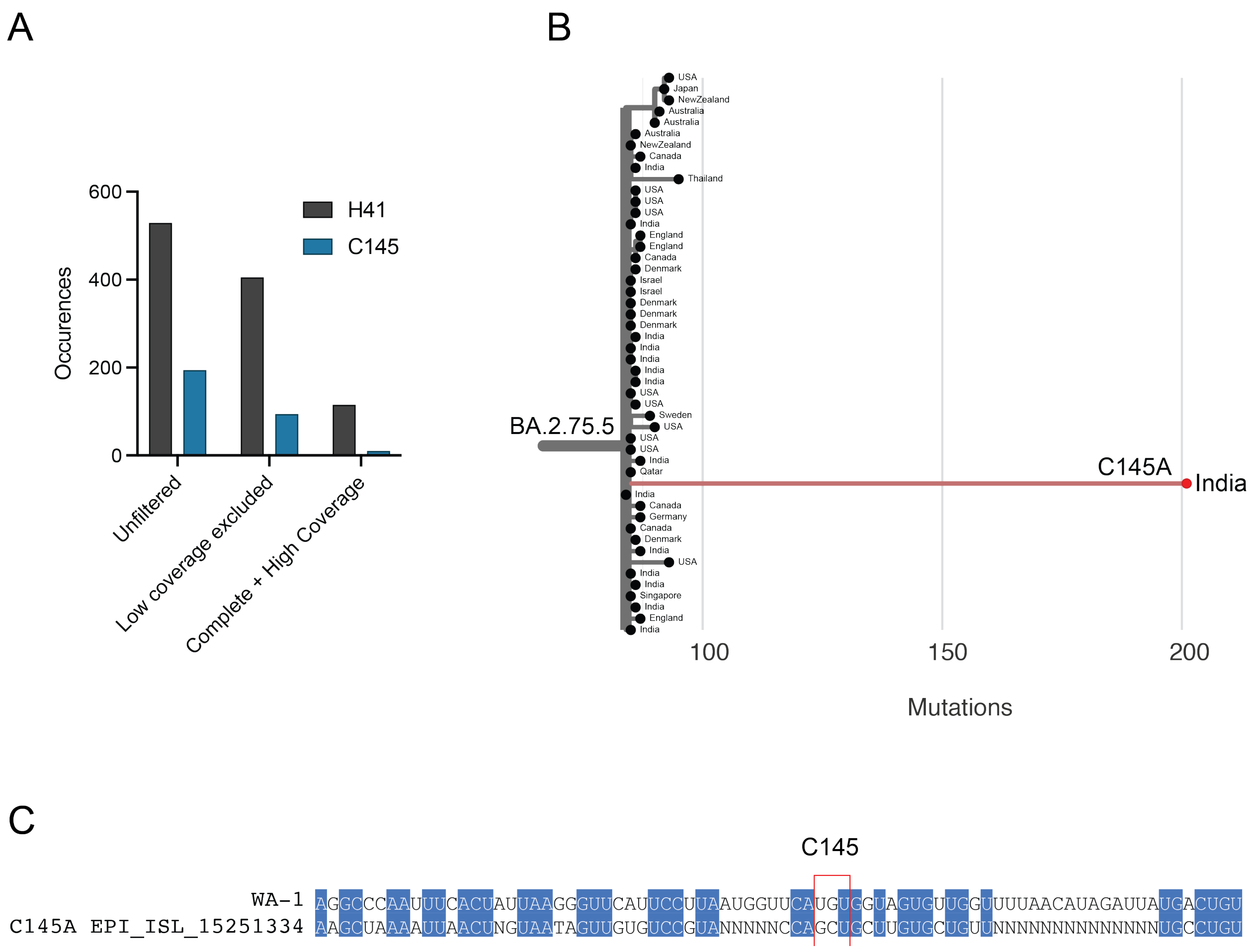


**Supplementary Figure 10. Mutations of the catalytic dyad found in GISAID.**
(**A**) The total number of amino acid changes reported for H41 and C145 in the GISAID database as of 5-Aug-2022 using the indicated filter settings in EpiCoV.
(**B**) Phylogenetic tree identifying closest common ancestor of a viral genome from India annotated with a C145A mutation in M$^{pro}$.
(**C**) M$^{pro}$ nucleotide sequence alignment of the C145 region of the ancestral WA-1 strain and of the mutant viral genome from India with C145A in panel (B).

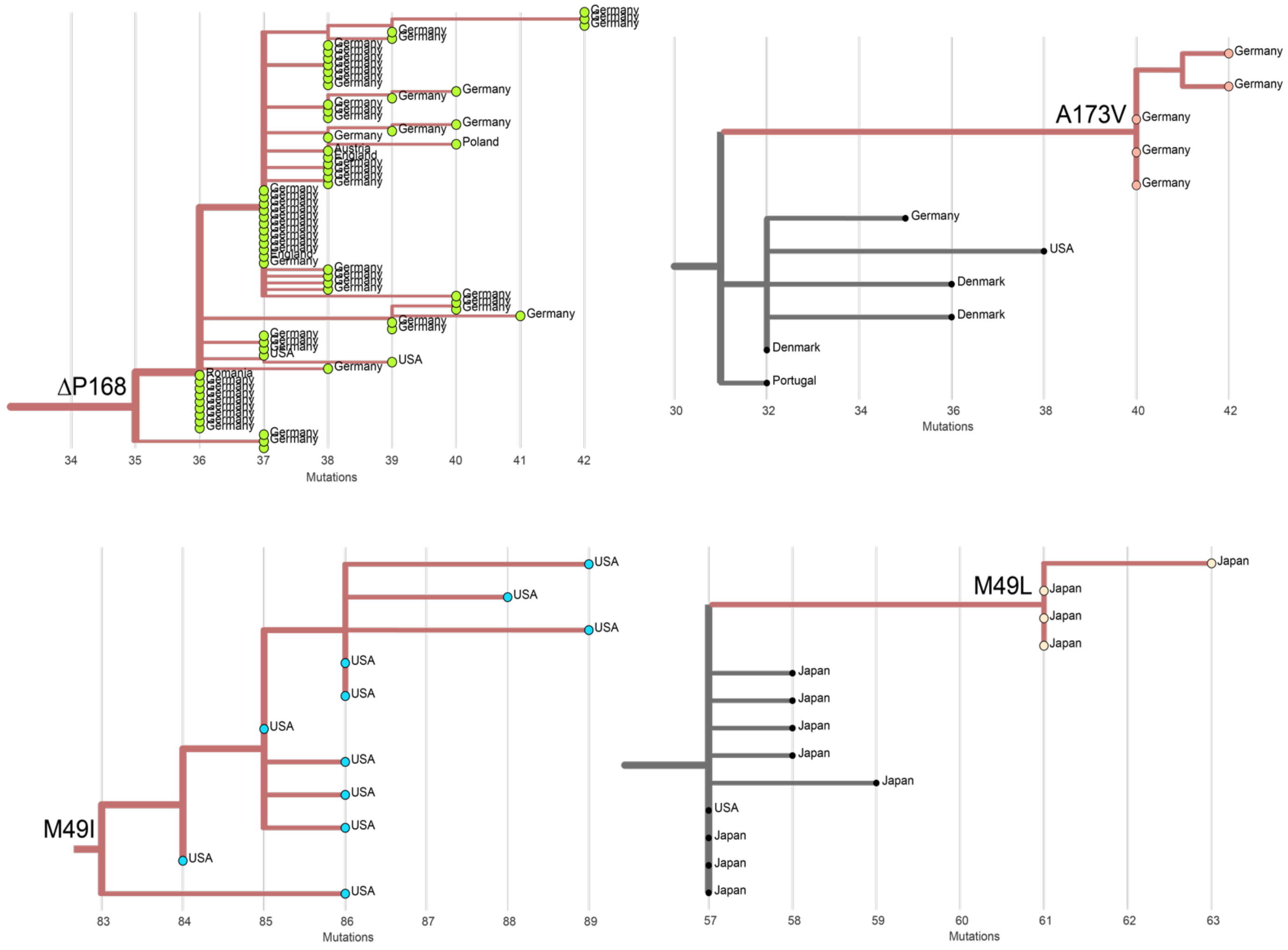


**Supplementary Figure 11. Examples of localized transmission of M^pro resistant variants.** Representative branches from phylogenetic trees in Fig. 5 blown-up to include geographic information and likely transmission chains.

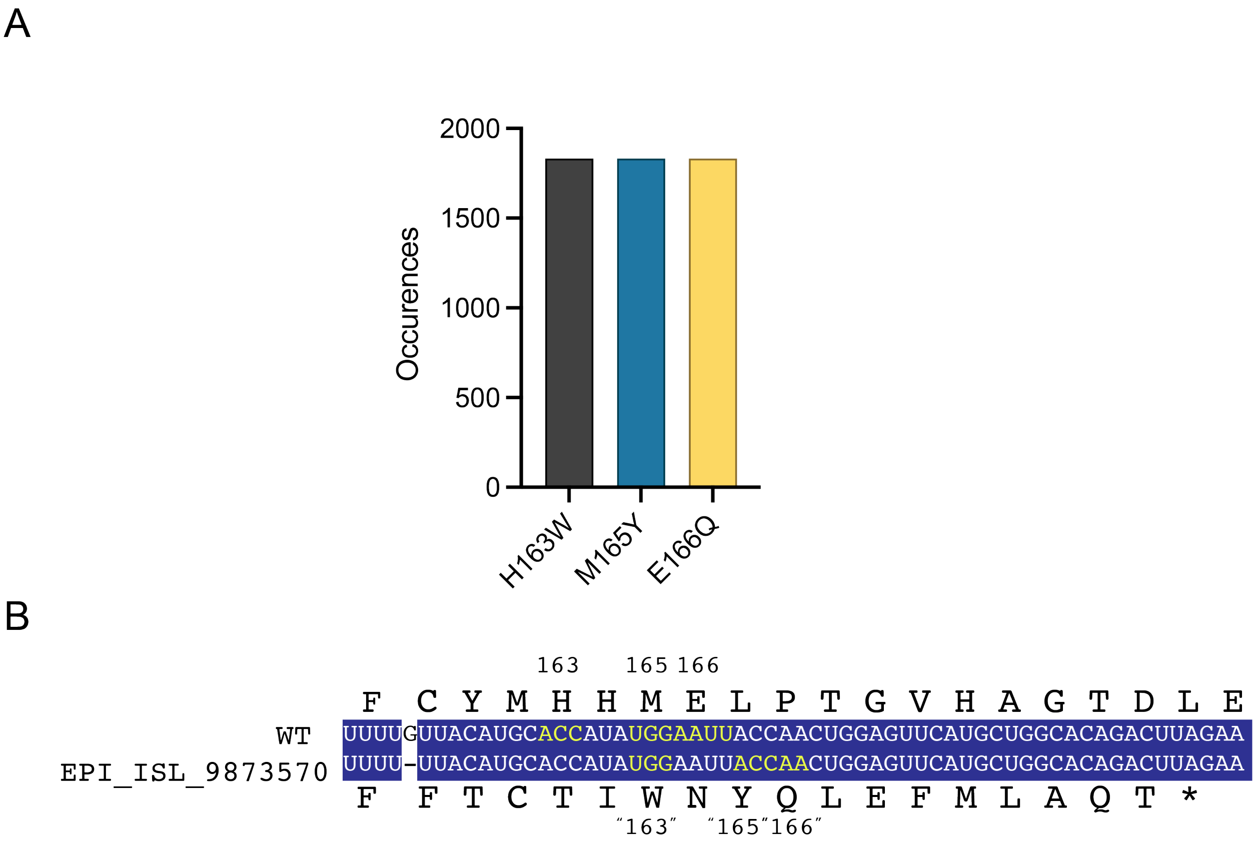


**Supplementary Figure 12. H163W, M165Y, and E166Q in the GISAID database are likely due to sequencing and annotation errors.**

(**A**) Curious co-occurrence of H163W, M165Y, and E166Q "mutants" in the GISAID database (31-July-2022).

(**B**) Alignment of WT M$^{pro}$ and a representative sequence from the GISAID database reported to harbor a deletion of G10533 (indicated by a "-") in a poly-U run. The resulting frameshift changes the coding sequence from M$^{pro}$ residue 160 onward including misalignments for "H163W", "M165Y", and "E166Q" (corresponding codons shaded yellow). It is not clear why the GISAID entry is annotated out-of-register (possibly because the artifactual frameshift is not considered in an automated data entry/annotation workflow). The reported sequence without the frameshift (*i.e.*, the corrected sequence) would be wildtype. We propose manual correction of these types of entries because some may contain true variants of interest.

**Table S1:** All protonation states for all titratable $M^{pro}$ residues considered in this work. Protonation states for aspartate, glutamate, lysine, cysteine, tyrosine, and arginine were determined with PROPKA3. Protonation states for histidine were determined by observed resolved protons on 7BB2's structure as well as from mechanistic interpretation (for residue His41). All protonation states were held consistent across all simulated $M^{pro}$ variants.

| Residue Name | Protonation State Assigned | Residue Numbers |
|---|---|---|
| Aspartate | Deprotonated | 33,34,48,56,92,153,155,176,187,197,216,229,245,248,263,289,295 |
| Glutamate | Deprotonated | 14,47,55,166,178,240,270,288,290 |
| Lysine | Protonated | 5,12,61,88,90,97,100,102,137,236,269 |
| Cysteine | Protonated | 16,22,38,44,85,117,128,145,156,160,265,300 |
| Tyrosine | Protonated | 37,54,101,118,126,154,161,182,209,237,239 |
| Arginine | Protonated | 4,40,60,76,105,131,188,217,222,279,298 |
| Histidine | Protonated ($N_\delta$) | 41,80,164 |
| Histidine | Protonated ($N_\varepsilon$) | 64,163,172,246 |

**Table S2:** System composition summary for each $M^{pro}$ variant constructed and simulated in this work.

| Variant | Box Size (Å x Å x Å) | Prot #Atoms | Wat #Atoms | Na, Cl #Atoms | Total #Atoms |
|---|---|---|---|---|---|
| WT | 90.7 x 97.2 x 114.4 | 9290 | 85389 | 88, 80 | 94847 |
| A173V | 90.7 x 97.2 x 114.4 | 9302 | 85392 | 88, 80 | 94862 |
| ΔP168 | 90.7 x 97.2 x 114.3 | 9262 | 85353 | 88, 80 | 94783 |
| DM | 90.7 x 97.2 x 114.3 | 9274 | 85344 | 88, 80 | 94786 |

**Table S3:** Summary of all molecular dynamics steps performed in this work. $^c$Indicates steps were performed with constraint on catalytic dyad residues. $^r$Indicates steps were performed with restraints on catalytic dyad residues. $^1$Indicates a timestep of 0.1 fs was used. $^2$Indicates a timestep of 0.2 fs was used. NAMD2.14 was used for all simulation steps except for NVT Pr (*i.e.*, the statistically relevant "production" sampling), in which NAMD3.0 GPU[12] was used.

| Variant | #Rep | Min (#step)$^c$ | Heat (ps)$^{c,1}$ | NpT Eq (ps)$^{r,1}$ | NVT Eq (ns)$^1$ | NVT Pr (ns)$^2$ |
|---|---|---|---|---|---|---|
| WT | 3 | 10,000 | 140 | 252 | 27.5 | 550 |
| A173V | 3 | 10,000 | 140 | 252 | 27.5 | 550 |
| ΔP168 | 3 | 10,000 | 140 | 252 | 27.5 | 550 |
| DM | 3 | 10,000 | 140 | 252 | 27.5 | 550 |
